# Structural basis for the intestinal protocadherin-based intermicrovillar adhesion complex

**DOI:** 10.64898/2026.05.11.724279

**Authors:** Akinobu Senoo, Pablo A. Guillen-Poza, Karin Fujishima, Hirofumi Kosuge, Takamasa Doumoto, Keisuke Kasahara, Tomohito Tanihara, Yuya Yoshida, Saeko Yanaka, Makoto Nakakido, Satoru Nagatoishi, Katsumi Maenaka, Shigehiro Ohdo, Naoya Matsunaga, Ruben Hervas, Kouhei Tsumoto, Jose M.M. Caaveiro

**Affiliations:** Laboratory of Protein Drug Discovery, Graduate School of Pharmaceutical Sciences, Kyushu University, 3-1-1 Maidashi, Higashi-ku, Fukuoka 812-8582, Japan; School of Biomedical Sciences, The University of Hong Kong, Hong Kong HKSAR, China; Department of Bioengineering, School of Engineering, The University of Tokyo, 7-3-1 Hongo, Bunkyo-ku, Tokyo 113-8656, Japan; Laboratory of Pharmacokinetics, Faculty of Pharmaceutical Sciences, Kyushu University, 3-1-1 Maidashi, Higashi-ku, Fukuoka 812-8582, Japan; Materials and Structures Laboratory, Institute of Integrative Research, Institute of Science Tokyo, 4259 Nagatsuta-cho, Midori-ku, Yokohama, Kanagawa 226-8503, Japan; Laboratory of Biomolecular Science, Faculty of Pharmaceutical Sciences, Hokkaido University, Sapporo, Japan; Medical Device Development and Regulation Research Center, School of Engineering, The University of Tokyo, 7-3-1 Hongo, Bunkyo-ku, Tokyo 113-8656, Japan; School of Biomedical Engineering, The University of Hong Kong HKSAR, China; Department of Chemistry and Biotechnology, School of Engineering, The University of Tokyo, 7-3-1 Hongo, Bunkyo-ku, Tokyo 113-8656, Japan

## Abstract

The intestinal brush border (BB), composed of densely packed microvilli on enterocytes, is essential for nutrient absorption and host defense. Its organization relies on the intermicrovillar adhesion complex (IMAC), mediated by protocadherins CDHR2 and CDHR5. Despite their clinical relevance in inflammatory bowel disease and several carcinomas, structural details of IMAC assemblies have remained elusive. Herein, we report the Cryo-EM structure of the adhesive complex at 3.4 Å resolution, revealing a heterotetrameric ensemble composed of a dimer of CDHR2 and a dimer of CDHR5. This assembly ensures uniform adhesive strength between neighboring microvilli, and facilitates hexagonal packing of microvilli. Biophysical analyses and molecular dynamics simulations revealed a kinked, Ca²⁺-free linker between domains EC3 and EC4 of CDHR5 conferring the necessary flexibility to withstand the shear stress caused during intestinal peristalsis. Collectively, these findings provide a structural framework for understanding BB organization and suggest strategies for therapeutics targeting IMAC in intestinal disorders.

## Introduction

The intestinal luminal surface is a frontline that is involved in the exertion of multiple essential functions such as nutrient processing, absorption(1, 2) and host defense against pathogenic bacteria(3–5); and thus is a tissue of crucial importance. In this context, enterocytes constitute one of the predominant cell populations. The apical surface of epithelial enterocytes lining the intestinal villi possesses a large number of densely packed protrusions termed microvilli (Figure 1A). Microvilli form bundles of themselves called brush border (BB) which serves as the site where the diverse functions of the intestine are executed. Microscopic observation of differentiated enterocytes reveals that microvilli within the BB are arranged in a close-packed configuration, exhibiting a geometrical “hexagonal array” configuration when observed from above(6, 7) (Figure 1A). Such naturally occurring high-ordered dense packing increases the apical surface area of the intestinal lining, thereby maximizing nutrient absorption, while simultaneously supporting a robust host-defense function.

**Figure 1.**
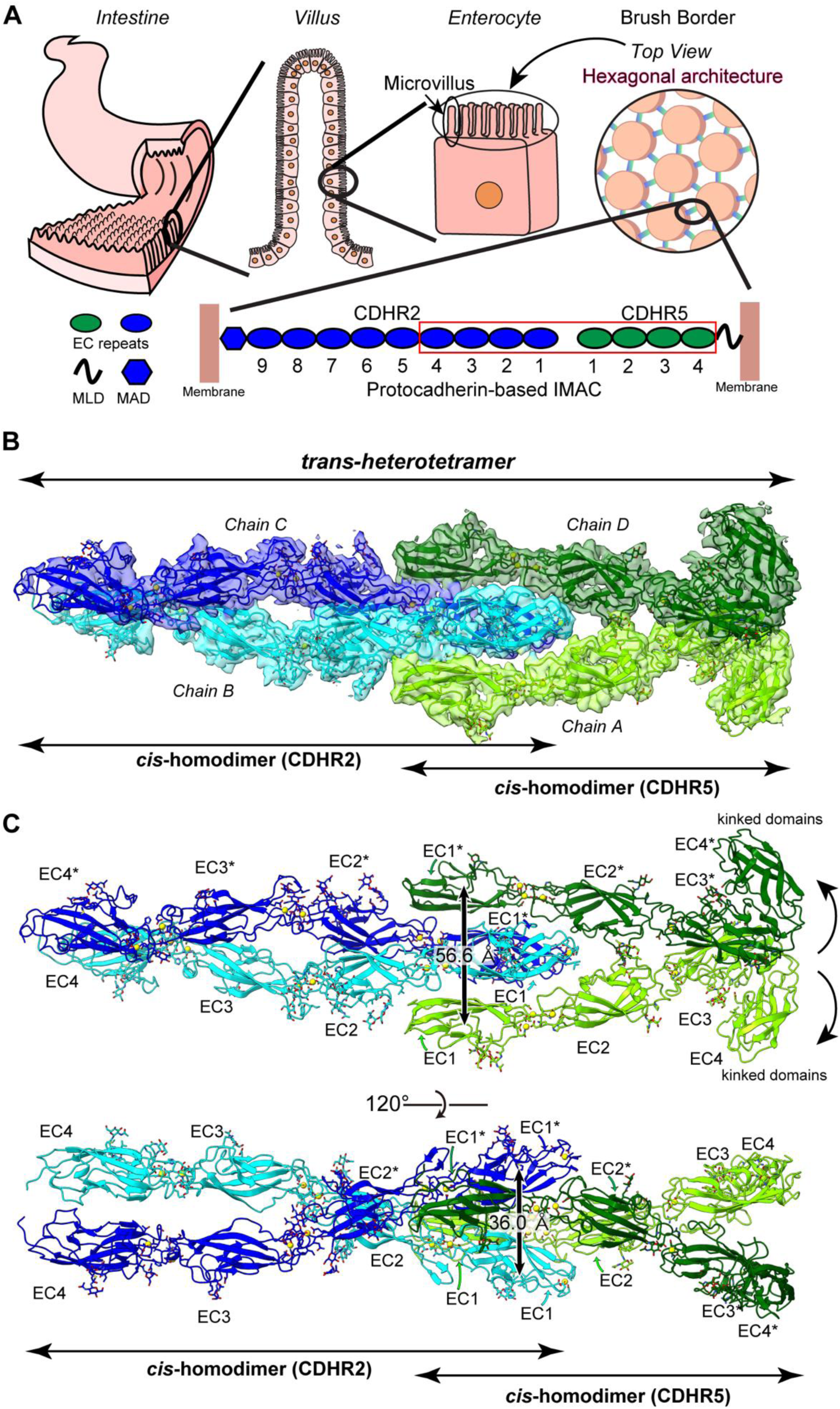
Molecular architecture of protocadherin-based IMAC. **(A)** Schematic illustration of hierarchical tissue structure in intestine. Brush border (BB) is formed as bundle of microvilli on the enterocytes, which manifests as hexagonal pattern as seen from the top view. Two types of protocadherin called CDHR2 and CDHR5 form the inter microvillar adhesion complex between microvilli. CDHR2 has nine EC repeats and a membrane adjacent domain (MAD), while CDHR5 has four EC repeats and a mucin-like domain (MLD). The red box shows the region of EC domains employed to solve the structure reported in this study. **(B)** Cryo-EM map of *trans*-heterotetramer and modeled structure in the map. Two chains of CDHR5 (chain A and chain D) are colored in light green and green; two chains of CDHR2 (chain B and chain C) are colored in cyan and blue. We named the interaction mode “crocodile-biting configuration” (see Supplementary Figure S3D for more information). **(C)** Built model of CDHR2 and CDHR5. EC1-4/EC1-4* from both CDHR2 and CDHR5 were being modeled. Glycosylation is shown in stick. Oxygen, nitrogen or calcium atoms are colored in red, blue or yellow, respectively. The unique kinked domain in EC3-4/EC3-4* of CDHR5 is shown by curved arrows. The distance between a mass center of EC1 and that of EC1* from CDHR5 is depicted with a black arrow. **(D)** Built model from another angle. The distance between a mass center of EC1 and that of EC1* from CDHR2 is depicted with a black arrow.

Formation of the BB is a process that entails bundling of adjacent microvilli, which critically depends on intermicrovillar adhesion mediated by two adhesion proteins(6). This adhesion module is referred to as the protocadherin-based intermicrovillar adhesion complex (IMAC). Classical biochemical analyses(8) and proteomics(9) have established that two adhesion proteins are localized to microvillar tips: CDHR2 (also known as PCDH24) and CDHR5 (also known as Mucin-like protocadherin or µ-Protocadherin). Knockout of either CDHR2 or CDHR5 results in defective brush-border assembly(10, 11). Besides, in patients with ulcerative colitis (UC), a type of inflammatory bowel disease, the expression level of CDHR5 is significantly reduced. Also, mice with low CDHR5 expression exhibit impaired brush-border formation accompanied by diminished host defense against microbes(11). These findings strongly suggest a close relationship between diseases and the role of IMAC formed by the two types of protocadherins. Emerging reports also link CDHR5 expressions to different cancer types such as colorectal cancer, renal cell carcinoma or pancreatic ductal adenocarcinoma(12, 13), underscoring its clinical relevance.

At the molecular level, CDHR2 and CDHR5, similar to classical cadherins, consist of sequential immunoglobulin-like domains involved in Ca²⁺-dependent IMAC assembly (Figure 1A). CDHR2 possesses nine extracellular cadherin (EC) repeats and a membrane-adjacent domain (MAD), followed by a single-pass transmembrane segment and a cytoplasmic tail. CDHR5 comprises four EC repeats, a mucin-like domain (MLD), a single-pass transmembrane segment and a cytoplasmic region. To date, several studies have focused on the intracellular regions, addressing the question of how these proteins are delivered to the tip of microvilli (14–17). By contrast, structural information on their ectodomains and specially the adhesive complex was poorly understood. Before our study, the only reported structures to date correspond to human CDHR5 EC1–2, human CDHR2 EC1–2, and mouse CDHR2 EC1–3(18). Crucially, although homo interaction of CDHR2 has been previously suggested(18), the heterotypic interface required for IMAC assembly has yet to be defined. Especially, hexagonal packing of microvilli requires uniform adhesive strength across all nearest-neighbor directions; otherwise, anisotropic forces would disrupt the carefully balanced hexagonal geometry. But how uniform adhesive strength originates from two different types of protocadherin proteins with different lengths is unknown. Besides, these proteins must achieve it in the presence of shear stress trying to break the IMAC caused by intestinal peristaltic pump(19, 20). The molecular features for the proteins to work under such a condition have not been elucidated either.

Herein, we report the cryogenic electron microscopy (cryo-EM) structure of the complex between CDHR2 EC1–4 and CDHR5 EC1–4. The adhesive complex is a supramolecular structure made of a *cis-*dimer of CDHR2 (the prefix *cis-* indicating that both protein units are located in the same microvillus) and a *cis*-dimer of CDHR5 that assemble in a *trans-*heterotetramer (where the prefix *trans*- indicates that each dimer is located on a different microvillus). Furthermore, we report a novel Ca²⁺-free linker between domains EC3 and EC4 of CDHR5 which adopted a kinked, flexible conformation. Through a combination of structural, biophysical, and computational analyses, we describe the structural features of the adhesive complex, illuminating the molecular basis allowing hexagonal packing in BB and adaptive strategies of these two types of protocadherin tailored to the environment of the intestinal milieu.

## Results

### Molecular architecture of an inter-microvilli adhesion complex

We determined the molecular architecture of protocadherin-based IMAC using cryo-EM with the optimized constructs of CDHR2 EC1-4 and CDHR5 EC1-4 (Supplementary Fig. S1, Table S1). After data processing, we determined the map to an average resolution of 3.41 Å. Data analysis and statistics are outlined in Supplementary Figure S2 and Table S2. From the density map, we were able to model all EC domains included in the constructs, i.e. CDHR2 EC1-4, EC1-4* and CDHR5 EC1-4, EC1-4*, where the asterisk indicates the domains of the second chain in each *cis-*dimer (Figure 1B). Local resolution of map regions corresponding to EC4/EC4* of both CDHR2 and CDHR5, as well as portions of EC1/EC1* of CDHR2 presented comparatively limited resolution, suggesting that these domains are dynamic in nature (Supplementary Figure S3A). With respect to the glycosylation of the protein, all putative N-glycan sites estimated by Uniprot, except for Asn405 of CDHR5, showed partial glycan occupancy that allowed model buildings (Supplementary Figure S4). In addition, glycosylation sites are distributed throughout all EC repeats, revealing that the protein is surrounded by an extensive glycan shield.

Model building of the structure revealed that both CDHR2 and CDHR5 formed a *cis*-homodimer (where *cis-* indicates that the protein chains comprising the homodimer are anchored in the same microvillus). The IMAC architecture is the result of the dimerization of a homodimer of CDHR2 anchored in one microvillus and a homodimer of CDHR5 anchored in an adjacent microvillus, resulting in a *trans*-heterotetramer bridging the two microvilli involved. We termed the *trans*-heterotetramer as the “crocodile-bite” configuration, because the widely open EC1-2/EC1-2* dimer of CDHR5 resembles the open jaws of crocodile capturing the EC1-2/EC1-2* dimer of CDHR2 (Figure 1B, Supplementary Figure S3B).

When focusing on the complex quaternary structure, we first noticed that the homodimer of CDHR5 showed widely open EC1-2/EC1-2* domains (56.6 Å between the center of mass of EC1 and that of EC1*). The homodimerization interface of CDHR5 involved the EC3-EC3* region with a calculated buried surface area (BSA) of 653.5 Å^2^ using the PDB PISA module(21) (Figure 1C, Supplementary Figure S3C, Table S3). This value is surprisingly small, given that the general protein-protein interaction surfaces cover more than 1,000 Å^2^(22); however, the nature of the dimer of CDHR5 EC1-4 is not questionable since it was robustly demonstrated by size exclusion chromatography with multi-angle light scattering (SEC-MALS) (Supplementary Figure S1). In addition to this, it is noteworthy to mention that a highly kinked conformation was observed in EC3-4 domains (Figure 1C), which is expected to play an important role in the dynamics of the protein (see below).

In contrast, the homodimer of CDHR2 showed a tighter packing arrangement of the EC1/EC1* domains (the distance between the center of mass of EC1 and that of EC1* was 36.0 Å) in which partial contacts between portions of EC2-EC2*, EC3-EC3* and EC4-EC4* were observed (Figure 1C, Supplementary Figure S3D, Table S4). Although the BSA value resulting from the interaction of all these interfaces was larger (1054 Å^2^), each component was relatively small (the largest being 523 Å^2^ between EC2 and EC2*) (Supplementary Figure S3D, Table S4). This could explain why CDHR2 EC1-4 required to be linked to the Fc region of IgG1 antibody to assist its stable dimerization for structural determination. However, it is important to point out that the full extracellular region comprising domains EC1-9 and MAD (EC1-9MAD) appeared as a homodimer, validating the structural approach (Supplementary Figure S1).

Finally, the interface analysis of the *trans*-heterotetramer bridging two neighboring microvilli indicated that the average BSA value between the two hetero interfaces was 757 Å^2^/interface (1513 Å^2^/homodimer of CDHR2 and CDHR5) (Table S5), again revealing a narrow interaction interface per protein chain. We discuss these features in more detail in the following sections.

### Characterization of CDHR5 homodimerization

We first attempted to characterize the nature of the CDHR5 *cis*-dimer observed in the adhesion complex. Following the structure, the interaction surface area between CDHR5 chains is mediated by the contact between an extruded loop comprising residues Cys240 to Ala251 and the β2-β3 sheets (Figure 2A). Seven residues with significant ASA (accessible surface area, >20 Å^2^) and a BSA/ASA ratio greater than 0.5 in both chains were found at the interface, namely Phe242, Val247, Ile249 and Gln250 belonging to the connecting loop, and Leu324, Tyr335 and Val337 belonging to the β2-β3 sheet (Table S3). These residues are exposed to the solvent in the monomeric state but appear largely buried upon dimerization. Among these, six residues are of hydrophobic nature, suggesting that the main driving force of homodimerization is the hydrophobic effect. Only one hydrogen bond, between Tyr335 and Asp244, was observed, although higher resolution data and therefore more accurate side-chain placement may reveal a greater number of them. The non-polar nature of the contact interface is also clear when coloring the protein surface by level of hydrophobicity, revealing significant hydrophobic patches around the area of interest (Figure 2A).

**Figure 2.**
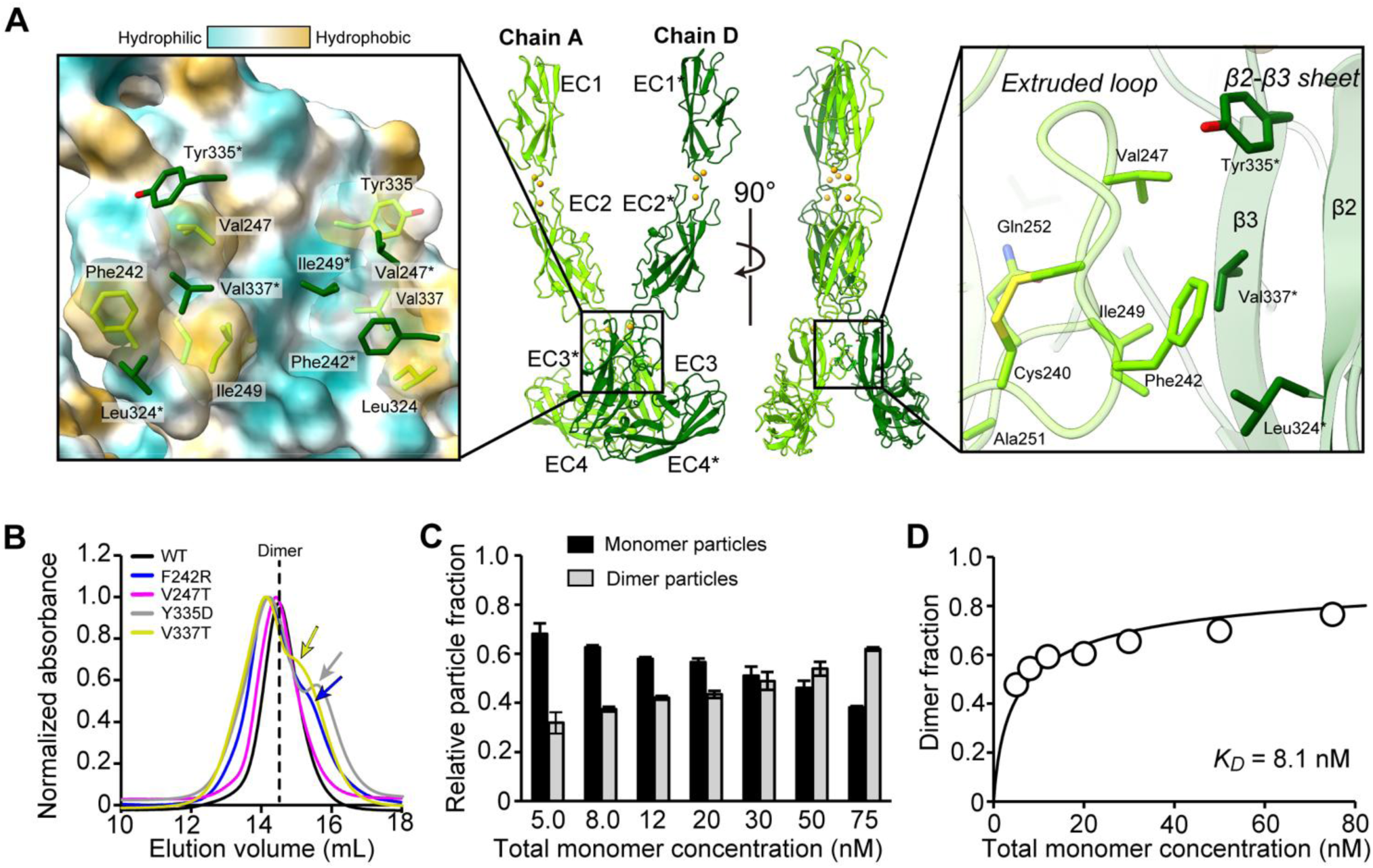
Characterization of homodimerization of CDHR5. **(A)** *Cis***-**homodimer of CDHR5 extracted from the complex structure from two different angles is shown in the middle panel. Chain A or D is colored in light green or deep green, respectively (same as Figure 1). Solid black boxes indicate the close-up regions on the right or left panels. The right panel depicts an enhanced view of interface. The interface of homodimerization consisted of an extruded loop from Cys240 to Ala251 of one chain and β2-β3 sheet from the other chain. Residues mentioned in the text are shown in sticks with oxygen, nitrogen or sulfur atoms with red, blue or yellow, respectively. The left panel shows a close-up view of the interface with surface colored by hydrophobicity using UCSF Chimera X software(47). One hydrophobic patch is made of Phe242, Val247, Ile249, and another of Leu324, Tyr335, Val337. On the surface representation from chain A, only side chain of residues from chain D is shown in stick. **(B)** SEC chromatograms from CDHR5 EC1-4 WT, F242R, V247T, Y335D and V337T in black, blue, purple, grey and yellow, respectively. The elution position of WT is shown by a dotted line. The peak height was normalized for the qualitative comparison. The colored arrows show the shouldered peaks for some mutants. **(C)** The monomer fraction and dimer fraction of counted particles measured by mass photometry at each concentration of total monomer. Black or grey bars show fraction of monomer or dimer, respectively. Error bars show standard deviation from three independent experiments. Raw data are shown in Supplementary Figure S5. **(D)** Plot of dimer fraction at each concentration of total monomer. The fitted curve is based on the least-square fitting.

From among the residues in the loop and β2-β3 sheet, the four residues Phe242, Val247, Tyr335, Val337 were selected for in depth analysis based on the high ASA/BSA ratio in both chains and contribution to hydrogen bond and subjected to mutational analysis. To profoundly change the nature of the molecular level interactions mediated by their side chains, these residues were mutated to display hydrophilic residues as follows: F242R, V247T, Y335D or V337T. SEC experiments were performed to qualitatively judge whether a change in their elution position/pattern occurs (or not). As a result, CDHR5 EC1-4 WT and its mutant V247T eluted as a monomodal peak at a position of 14.5 mL, while F242R, Y335D or V337T showed bimodal peaks at 14.3 and 15.3 mL (Figure 2B), indicating the emergence of species with smaller molecular size. These results indicate that the perturbation of the hydrophobic environment at the interface partially disrupted the dimeric nature of CDHR5 EC1-4.

To determine the dissociation constant (*K*_D_) of the CDHR5 EC1-4 dimer, we employed mass photometry, an interferometric scattering microscopy to determine the molecular weight of macromolecules (23). Following a previous study(24), the number of particles corresponding to dimers and monomers were quantified at several concentrations of the sample (Supplementary Figure S5, Figure 2C). As shown in Figure 2C, as the concentration of the CDHR5 increased, the fraction of monomer species decreased and simultaneously the fraction of dimer species increased, indicating that the monomer-dimer equilibrium exists around the concentration tested. When the fraction of dimers was plotted against the total concentration of monomers, a *K*_D_ value 8.1 nM was determined, indicating a strong affinity between the molecules of CDHR5 (Figure 2D).

### Characterization of the adhesive interface between CDHR2 and CDHR5

The interaction between CDHR5 and CDHR2 takes place mostly between their corresponding EC1 domains (Figure 3A). Indeed, the density map corresponding to the interface revealed that there is a vacant space between EC2 of CDHR5 and EC1 of CDHR2 (Figure 3A), consistent with the observation that the map density of EC1 of CDHR2 was weak (Supplementary Figure S3A). At the interface, there are several residues showing high ASA/BSA ratio (> 0.5) in both interfaces such as Ser90, Leu91, Val105, Thr106, Arg109, Phe111 and Ser113 (Figure 3B, Table S5). Especially, R109G is a mutation previously employed to reduce the formation of brush borders in Caco-2 bbe cells (6), and that our structure explains why it was effective at that. The density corresponding to the Arg109 and surrounding residues suggests that this arginine gives rise to cation-pi interactions with Tyr91 and Tyr87 of CDHR2, respectively (Figure 3C).

**Figure 3.**
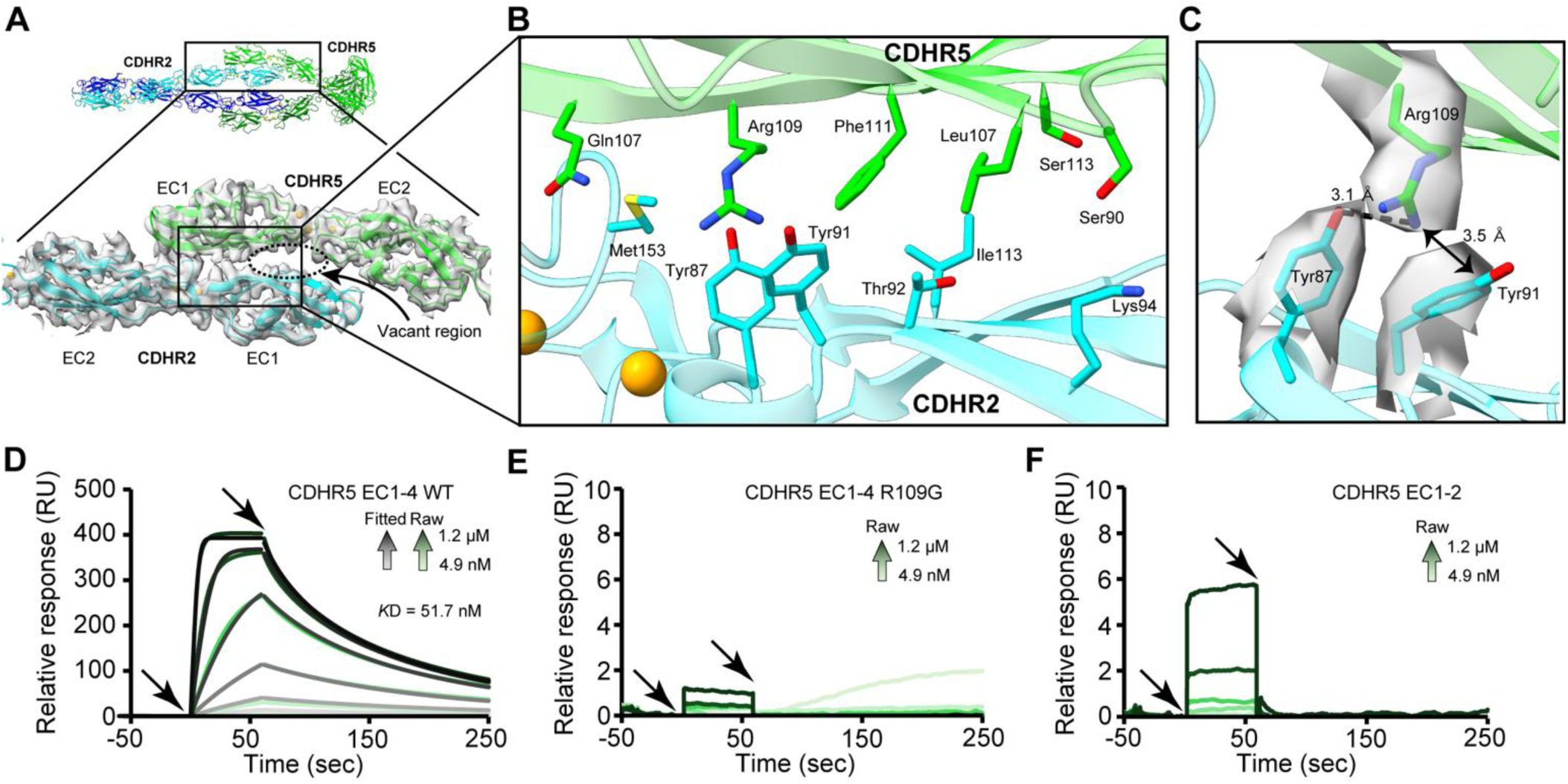
Characterization of hetero interaction. **(A)** Interface of hetero interaction. The interface is localized to EC1 of CDHR2 and CDHR5 shown in cyan or light green, respectively, with cryo-EM map of corresponding area. The arrow shows the space in the interface. **(B)** The enhanced view of hetero interface. Residues mentioned in the text are shown in stick with oxygen, nitrogen and sulfur atoms in red, blue or yellow, respectively. Calcium atoms are shown in orange spheres. **(C)** The enhanced view of Arg109 from CDHR5 and its interacting residues from CDHR2 (Tyr87 and Tyr91) with cryo-EM map of them. Arg109 likely forms cation-pi interaction with Tyr91 and a hydrogen bond with Tyr87. The distance between NH1 from Arg109 and CA from Tyr91, NH1 from Arg109 and OH from Tyr87 are depicted in black arrow or dotted line, respectively. **(D) (E) (F)** SPR sensorgrams of CDHR5 EC1-4 WT **(D)**, EC1-4 R109G **(E)** and EC1-2 WT **(F)** as an analyte. In all conditions, CDHR2 EC1-4-Fc was immobilized on Sensor Chip Protein A. The analyte solutions were adjusted to 1200, 400, 133, 44.4, 14.8 and 4.94 nM. The green traces show the raw data, while the black traces show the fitted data. For **(E)** and **(F)**, no fitting data is shown due to poor reliability. Note that scale of vertical axis is 50 times different from that of panel **(D)**.

To quantitatively analyze the effect of each residue on the formation of the adhesive complex, we employed surface plasmon resonance (SPR). The construct of CDHR2 EC1-4-Fc was immobilized on a Sensor Chip Protein A, whereas CDHR5 EC1-4 WT or its mutants were injected into the sensor chip surface at several different concentrations. We examined R109G, S90A and Q107A as examples of H-bond donor or acceptor. Also, we tested L91W to evaluate whether filling up the gap at the interface will affect the interaction. CDHR5 EC-14 WT showed fast association rate (*k*_on_ = 2.72×10^5^ M^−1^s^−1^) and slow dissociation rate (*k*_off_ = 1.42×10^−2^ s^−1^), resulting in apparent *K*_D_ value of 51.7 nM (Figure 3D, Table 1). The slow *k*_off_ value may be attributed to avidity effect of the *cis*-dimer of CDHR5 EC1-4. This *K*_D_ value is much stronger than that of classical cadherin (µM order) (25). The contribution of Arg109 proved to be critical, since the R109G mutant practically abolished the interaction between the cadherin moieties (Fig. 3E). In contrast, S90A or Q107A did not have quite an effect as strong as that in R109G, showing a meager 2.7- or 4.8-fold decrease in affinity (Supplementary Fig.S6A, S6B, Table 1). L91W showed slightly improved affinity with respect to the unmodified protein (apparent *K*_D_ = 42.2 nM), thus validating the contribution of a bulky side chain at this position (Supplementary Fig.S6C, Table 1).

**Table 1.** Kinetics parameters from SPR experiments.

| Protein | Oligomerization state | $k_{on} (M^{-1}s^{-1}) \times 10^{-5}$ | $k_{off} (s^{-1}) \times 10^2$ | $K_D$ (nM) | Affinity loss |
| --- | --- | --- | --- | --- | --- |
| EC1-4 WT | dimer | 2.7±0.009 | 1.4±0.004 | 51±0.2 | - |
| EC1-4 S90A | dimer | 2.9±0.008 | 4.0±0.01 | 140±0.5 | 2.7 |
| EC1-4 L91W | dimer | 4.4±0.02 | 1.9±0.008 | 42±0.3 | 0.82 |
| EC1-4 Q107A | dimer | 3.6±0.02 | 9.0±0.04 | 250±2 | 4.8 |
| EC1-4 R109G | dimer | N.D. | N.D. | N.D. | > 1000-fold |
| EC1-4 Y335D | monomer | N.D. | N.D. | N.D. | > 1000-fold |
| EC1-2 WT | monomer | N.D. | N.D. | N.D. | > 1000-fold |

N.D.: not determined because the binding response was too low or non-specific.
The errors are standard errors from global fitting of concentration series.

**Table 2.** Thermodynamic parameters from DSC experiments.

| Protein | Construct | Fitting 1 |  | Fitting 2 |  | Fitting 3 |  |
| --- | --- | --- | --- | --- | --- | --- | --- |
| | | $\Delta H$ (kcal/mol) | $T_m$ (°C) | $\Delta H$ (kcal/mol) | $T_m$ (°C) | $\Delta H$ (kcal/mol) | $T_m$ (°C) |
| CDHR5 | EC1-4 | 20.1 | 60.0 | 41.0 | 65.6 | 41.9 | 74.9 |
| CDHR5 | EC1-2 | 130 | 74.23 | N.A. | N.A. | N.A. | N.A. |
| CDHR2 | EC1-4 | 111 | 82.2 | N.A. | N.A. | N.A. | N.A. |
| CDHR2 | EC1-2 | 37.8 | 69.8 | 60.5 | 75.3 | N.A. | N.A. |

N.A.: not applicable

As shown in Supplementary Figure S1, EC1-4 of CDHR2, which is shown to be monomeric, did not interact with CDHR5 EC1-4, indicating that the monovalent interaction between CDHR2 and CDHR5 is weak. To elaborate on this idea, EC1-2 or the latter fraction of EC1-4 Y335D were injected into the sensor chip previously decorated with CDHR2 EC1-4-Fc. Both EC1-2 and the mutant EC1-4 Y335D retained the hetero (*trans-*)interacting domains, but in a monomeric protein that cannot form the *cis*-dimer. As shown in Figure 3F and Supplementary Figure S6D, the binding response from both analytes drastically decreased, practically to zero, compared with the WT construct of EC1-4 (homodimer). These results support the idea of significantly weak hetero interaction in monovalent form.

### Dynamic nature of the kinked domains of CDHR5

We next attempted to examine the kinked domains (EC3-4 domains) of CDHR5. One important aspect of the kinked domains is their potential flexibility, which is connected to their biological relevance. Flexibility is often attributed to Ca^2+^-free interdomain linkers or a linker with less than three Ca^2+^ ions. For example, in the case of PCDH15, another class of cadherin, the Ca^2+^-free linker is thought to contribute to its mechano-sensing function by its flexible nature (26–28). Therefore, we first investigated the number of Ca^2+^ ions at each interdomain linker in CDHR5.

The canonical three Ca^2+^ binding motifs consist of XEX and DXE consensus sequences at the preceding EC repeat, DXNDN at the interdomain linker, and DXD and XDX from the following EC repeat(29). When the sequence of CDHR5 is analyzed, a few incomplete or missing motifs were identified, such as DXE and DXNDN in EC2, and XEX, DXE, XDX and DXNDN in EC3 (Figure 4A). As reported in EC1-2 structure of CDHR5(18), clear density was observed in canonical motifs around the bottom of EC1 and top of EC2 in which three Ca^2+^ ions can be modeled (Figure 4B). Similarly, there was a clear density to support the Ca^2+^-binding at the top of the EC3 region (Figure 4C). However, the density around the bottom of EC2 did not support the modelling of two of the three Ca^2+^ ions, nor that around the bottom of EC3 and the top of EC4, presumably because of the high incompletion of the motifs as stated above (Figure 4D). Taken together, CDHR5 displays one Ca^2+^ atom between EC2 and EC3, and a Ca^2+^-free linker between EC3 and EC4 as summarized in Figure 4E. Therefore, sequence analysis and experimental data coincide to predict the same Ca^2+^ ion distribution, which had also been predicted by AlphaFold3 (Supplementary Figure S7). Note that unlike CDHR5, almost all the Ca^2+^ binding motifs were conserved in CDHR2 EC1-EC9, suggesting that CDHR2 has relatively rigid and straight conformation (Supplementary Figure S7), although we could not model Ca^2+^ ions in some of the regions due to lower local resolution at these positions.

**Figure 4.**
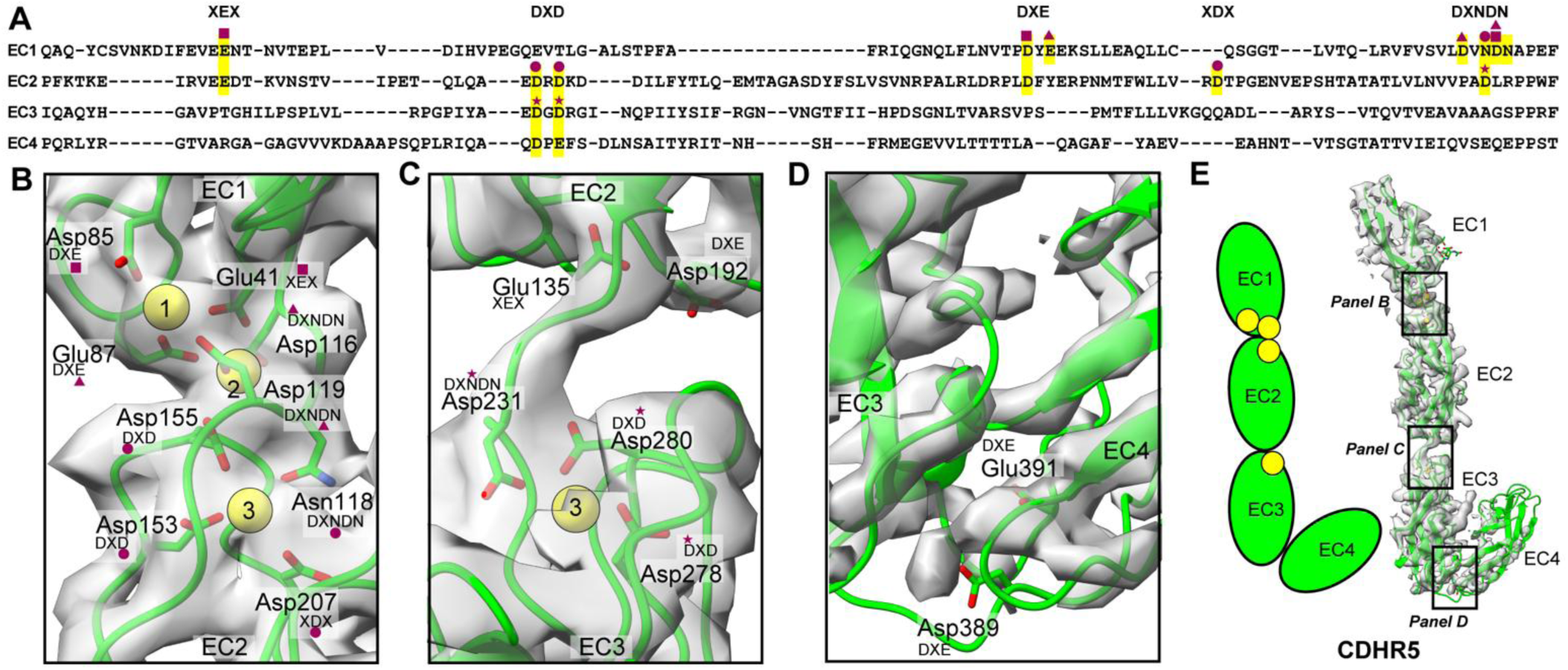
CDHR5 has Ca^2+^-free linker. **(A)** Primary sequence of CDHR5 (Gln26 to Thr459). Conserved canonical Ca^2+^-binding motifs are highlighted with yellow background. Also, the motifs (XEX, DXD, DXE, XDX and DXNDN) are highlighted on top of sequence. The residues coordinating to Ca^2+^ in the structure are marked with purple square, triangle, circle or star. **(B) (C) (D)** The Ca^2+^-binding region between EC1 and EC2 **(B)** or EC2-EC3 **(C)** or EC3-EC4 **(D)**. Cryo-EM density map is shown at the same counter level in grey. Residues are shown in sticks with oxygen, nitrogen or calcium atoms colored in red, blue or yellow, respectively. For each residue, the corresponding motif is added. If a residue is used to bind to Ca^2+^, the same mark as **(A)** (purple square, triangle, circle or star) are also added. **(E)** Reduced view of an extracted chain from CDHR5 homodimer with density map in grey. Each box shows the enhanced region in panel **(B)**∼**(D)**. The schematic illustration summarizes the conclusion of Ca^2+^-binding in each interdomain region.

Based on the Ca^2+^ modeling above, we evaluated the level of flexibility of the domains from CDHR5 using molecular dynamics (MD) simulations. To simplify the starting structure, we extracted EC1-2/EC1-2* from CDHR2 and placed it in complex with CDHR5 EC1-4/EC1-4* (Supplementary Figure S8A). At the glycosylated asparagine positions, we modeled agalactosylated biantennary N-glycan (G0 glycans) to the position where the density of a part of glycan was visible. Three independent runs of 100 ns were performed, and the convergence of the simulations was confirmed by RMSD values of the trajectories (Supplementary Figure S8B). The trajectories from 25 ns to 100 ns were used for the subsequent analysis. To understand the flexibility of the complex, we first computed the RMSD values of CDHR5 EC2/EC2*, EC3/EC3* or EC4/EC4* against CDHR2 EC1/EC1* as references. As shown in Figure 5A-5C, the averaged RMSD value were 9.6/8.5 Å, 25.1/18.4 Å or 29.6/38.6 Å for CDHR5 EC2/EC2*, EC3/EC3* or EC4/EC4*, respectively, becoming larger in the order of EC2 < EC3 < EC4. This trend seems to reflect the order of flexibility caused by the three-Ca^2+^-linker, the one-Ca^2+^-linker and the Ca^2+^-free linker, respectively. To visually show the displacement of each domain, spatial traces from the trajectories were depicted (Figure 5D-5F). As expected from the RMSD values, the volume of the spatial trace became largest in EC4. When the volume of each spatial trace was measured, volumes of 5.02×10^3^/1.05×10^4^ Å^3^, 4.04×10^4^/3.86×10^4^ Å^3^ or 6.71×10^4^/6.18×10^4^ Å^3^ for CDHR5 EC2/EC2*, EC3/EC3* or EC4/EC4*, respectively, were determined. In a representative snapshot from the trajectories, the angle calculated from the mass centers of EC3 or EC4 and the Cα of a residue from domain linker was 132.7°, corresponding to an almost straight conformation (Supplementary Figure S8C). When this snapshot was superposed with the starting structure, the structural change was not only the open conformation of EC3-EC4 angle but also reflected in the twisting movement seen from the top view (Supplementary Figure S8C). Overall, the MD simulations demonstrate that the possible covering space from CDHR5 EC3-4 is widely enhanced due to the Ca^2+^-free linker, indicating the flexible nature of these domains.

**Figure 5.**
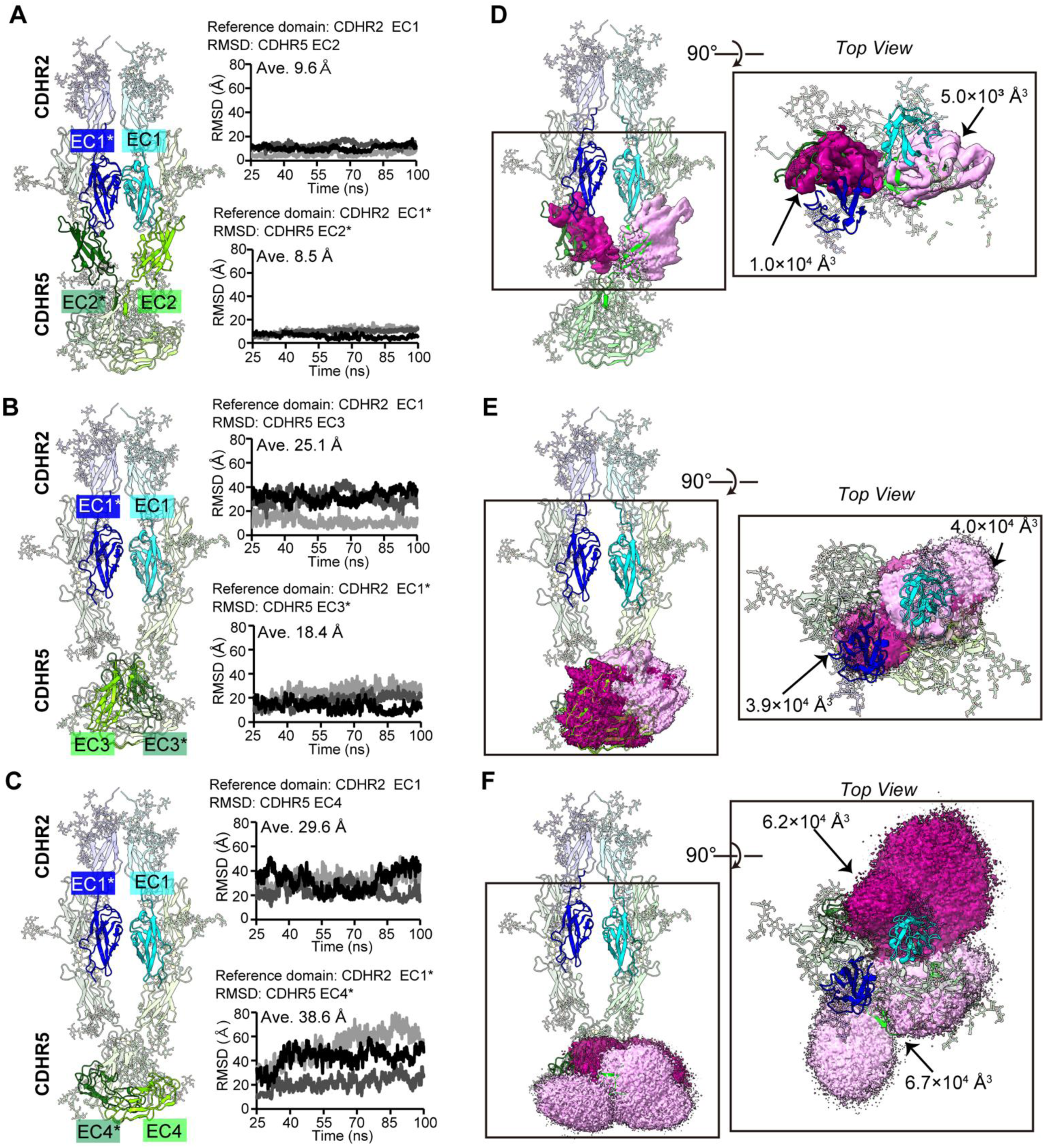
Dynamics of kinked EC3-EC4 domains in MD simulations. **(A) (B) (C)** RMSD values from MD trajectories of EC2/EC2* **(A)**, EC3/EC3* **(B)**, EC4/EC4* **(C)** of CDHR5 when EC1/EC1* of CDHR2 was used as a reference. Three independent trajectories from MD simulations correspond to black, grey and light grey. The average RMSD values are written. The left structures show EC1/EC1* of CDHR2 as references or each domain from CDHR5 to calculate RMSD in cartoons without transparency; the other domains are shown in transparent cartoons. We note that RMSD values of EC1/EC1* of CDHR5 with EC1/EC1* of CDHR2 used as a reference was small enough (4.7 Å/3.7 Å), respectively. **(D) (E) (F)** The spatial trace (moving region) of EC2/EC2* **(D)**, EC3/EC3* **(E)**, EC4/EC4* **(F)** of CDHR5 against EC1/EC1* of CDHR2 are depicted in maps in pink/magenta. Spatial trace maps generated from MD simulations showed different maximum density values among systems. To allow direct visual comparison, all maps were displayed at isosurface levels corresponding to 10% of their respective maximum density values. The measured volume of each spatial trace is written.

### Stability of CDHR2 and CDHR5

Proteins exposed to shear stress of microvilli caused by peristaltic pumping or by bacterial infections necessitate strong molecular stability. To understand the robustness of the adhesive complex as well as their protein subunits, we first evaluated the thermostability of CDHR2 and CDHR5 using differential scanning calorimetry (DSC). Three peaks were detected from the fitting of the raw data obtained with CDHR5 EC1-4 (Figure 6A). The major transition midpoint (also known as melting temperature or *T*m) of the peaks were 65.6 ℃ and 74.9 ℃, and a minor pre-transition at 60 ℃. Because CDHR5 EC1-2 showed a single peak of *T*m = 74.2 ℃ (Figure 6B), the two peaks with lower *T*m from CDHR5 EC1-4 seem to be derived from EC3 or EC4 connected with Ca^2+^-deficient linkers. Thermostability of CDHR2 proved to be higher than CDHR5, with *T*m values of 82.2 ℃ for EC1-4, or 69.8 ℃/75.3 ℃ for EC1-2 (Figure 6C, 6D). We also evaluated the thermostability of the CDHR5 mutants prepared above. To this end, we employed a high-throughput technique called differential scanning fluorimetry (DSF). We first evaluated CDHR5 EC1-4 WT. As shown in Figure S9 and Table S6, two clear peaks were observed at 72.2 ℃ or 63.2 ℃, indicating that the trend from DSF analysis is comparable to that of DSC with slightly lower *T*m values. Based on Figure 6A and 6B, it is reasonable to consider that these two peaks correspond to the denaturing of EC1-2 and EC3-4, respectively. Subsequently, the mutants from homodimerization interface (F242R, V247T, Y335D and V337T) were evaluated in DSF. Interestingly, the peak with lower *T*m value corresponding to the unfolding event of EC3-4 was barely seen for all mutants, suggesting that EC3-4 was destabilized by the mutations to the level in which cooperative unfolding event was undetectable (Figure S9A, Table S6). Since these residues seem to be important for the complete establishment of homodimerization, the inhibition of homodimerization seems to destabilize the local folding in EC3-4, although the overall tertiary structure of EC1-4 will not be largely affected by the mutations based on the similar main elution volumes of SEC among mutants in Figure 2B. On the contrary, mutants tested in hetero interaction analysis (S90A, L91W, Q107A and R109G) retained two clear peaks, showing that these mutations did not affect the stability of EC1-4 (Figure S9B, Table S6).

**Figure 6.**
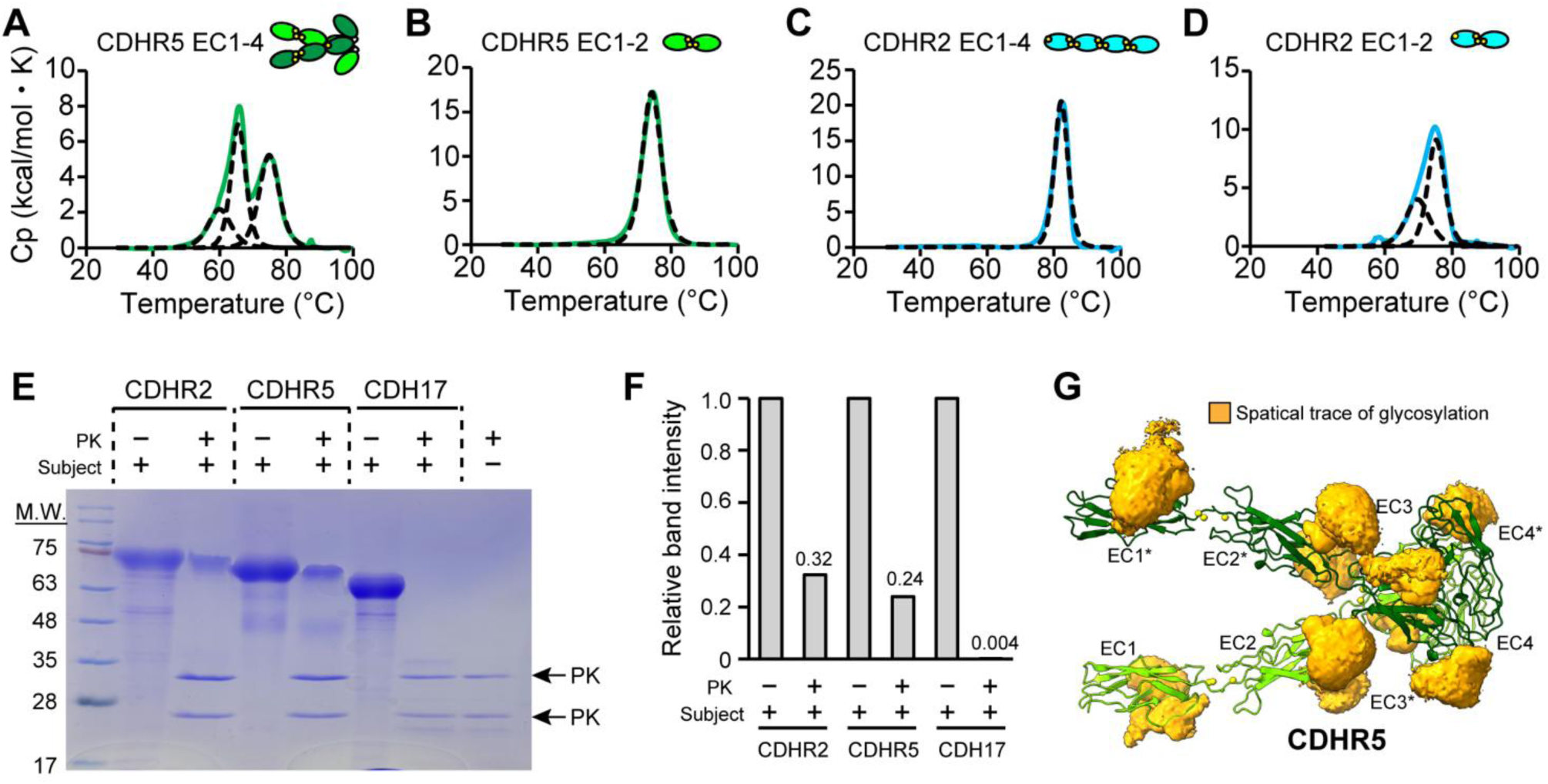
Molecular stability of CDHR2 and CDHR5. **(A)∼(D)** DSC analysis of CDHR5 EC1-4 **(A)**, CDHR5 EC1-2 **(B)**, CDHR2 EC1-4 **(C)** and CDHR2 EC1-2 **(D)**. Schematic illustration shows the domains and oligomerization state of samples. In each panel, raw data are shown in green or cyan solid lines, while fitting data are shown in black dotted lines. **(E)** Protein digestion assay using Proteinase K (PK). Each subject sample (CDHR2 EC1-4, CDHR5 EC1-4, CDH17 EC1-4) were incubated with PK and evaluated in SDS-PAGE. **(F)** Quantification of the band intensity using Image J software. The intensities of the bands with or without PK for each subject were quantified. On top of each bar, the actual values were written. For each protein, the band intensity of a subject without PK was normalized to 1.0. **(G)** Spatial traces of N-linked glycans on CDHR5 from MD simulations. All maps were displayed at isosurface levels corresponding to 5% of their respective maximum density values.

The high thermostability drove us to test the proteinase resistance of CDHR2 and CDHR5. Since CDHR2 and CDHR5 are constantly exposed to proteases secreted by bacteria that try to enter the enterocyte, CDHR2 and CDHR5 may utilize their rigidity to tolerate the degradation by protease. To investigate this, we performed protein digestion assay using Proteinase K (PK). PK is a non-specific protease that can digest proteins randomly. The cadherin protein CDH17, also expressed in intestinal cells and involved in adherence junctions (but not stabilizing microvilli) was employed as a comparison. Protein samples of CDHR5 EC1-4, CDHR2 EC1-4, or CDH17 EC1-4 were mixed with PK and the results were examined with SDS-PAGE. As shown in Figure 6E and 6F, the bands corresponding to the whole protein of CDHR5 and CDHR2 EC1-4 were detected, while that of CDH17 EC1-4 was not. The quantification of band intensity indicated that 32% of CDHR5 and 24% of CDHR2 remained intact in the presence of PK (Figure 6F). These results demonstrate that CDHR2 and CDHR5 acquired higher protease resistance compared to the cadherins responsible for adherence junction, consistent with the order of thermostability.

## Discussion

In this study, we report on the molecular architecture of protocadherin-based intermicrovillar adhesion complex (IMAC) determined by cryo-EM and elucidated its assembly mechanism using various biophysical and computational techniques. The results pave the way to answer the questions raised in the introduction; (1) how uniform adhesive strength required for the hexagonal packing of microvilli originates from two different types of protocadherin proteins and (2) how the proteins work in the presence of shear stress caused by peristaltic pumping (and gastrointestinal motility in general).

The answer to question (1) is a strict control to establish hetero interaction in a single species of complex. Small hetero interaction surface area per chain helps suppress monomer–monomer hetero interactions; instead, robust IMAC assembly occurs only between homodimers engaging at a correct angle. Also, SEC-MALS experiments did not show the peak of *trans*-homotetramer (i.e. association of two microvillus by two CDHR2 homodimers or by two CDHR5 homodimers), suggesting that *trans*-homo interaction is very weak or non-existing. In this way, the molecular complex species generated from two protocadherin proteins is limited to homogeneous adhesion distance and strength (Figure 7).

**Figure 7.**
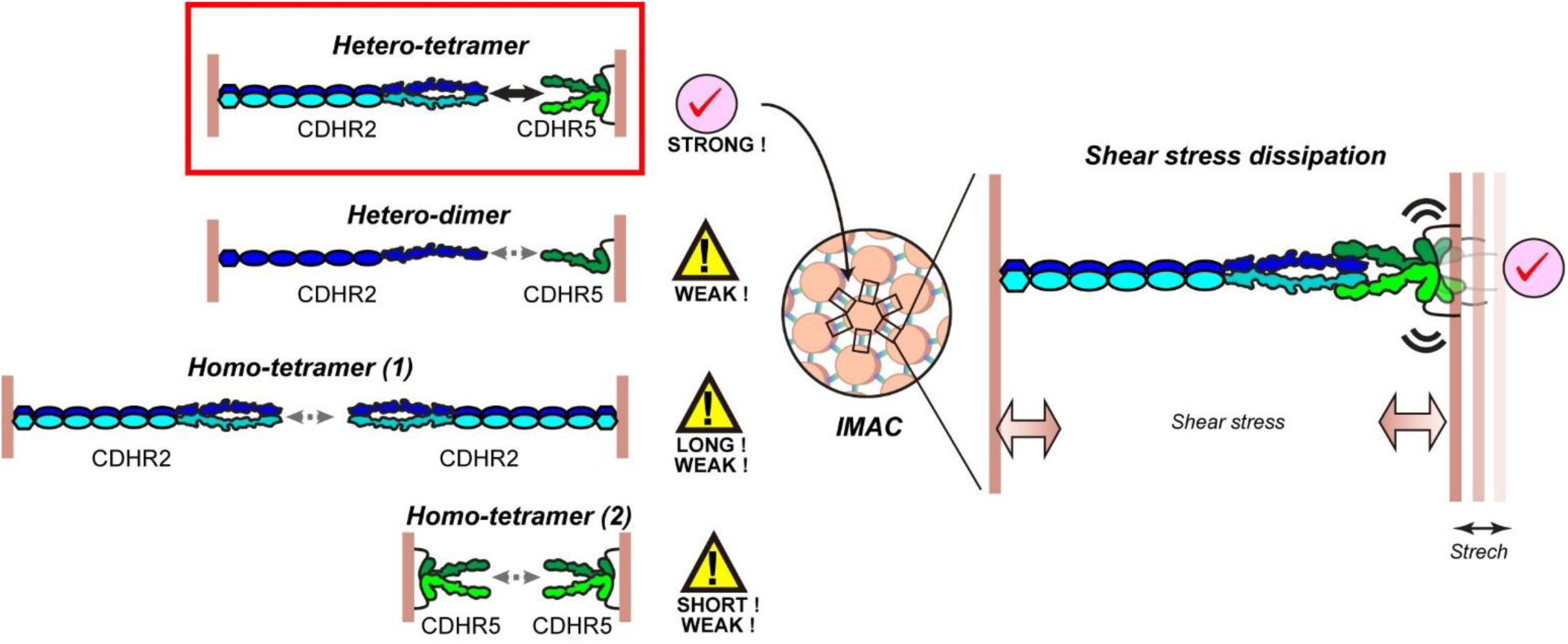
Model of IMAC formation. This study revealed that only strong interaction occurs between homodimers of both CDHR2 and CDHR5 (i.e. hetero-tetramer bridging the neighboring microvilli). The hetero interaction mediated by monomer CDHR2 and CDHR5 or *trans*-homo interaction (homo-tetramer (1) and homo-tetramer (2) bridging neighboring microvilli) by only CDHR2 or CDHR5 will be weak enough. This strict control for the establishment of bridging microvilli will thus keep the length and strength among microvilli uniform, contributing to the formation of hexagonal architecture of brush borders. Once the hetero interaction is formed, its strength is strong. In addition, the Ca^2+^-free linker at EC3-4 domains of CDHR5 allows the kinked conformation and flexible dynamics of these domains. This flexibility will help to dump the shear stress arising from intestinal peristalsis. Collectively, the molecular architecture achieves the balanced hard and soft required in the circumstances on the surface of intestine.

By operating in this manner, the system also seems to suppress hetero interactions between CDHR2 and CDHR5 on a single microvillus, thereby allocating these molecules efficiently to intermicrovillar adhesion. This will be achieved by two mechanisms. First, it would be difficult for the more rigid CDHR2 to sufficiently bend on itself to face a CDHR5 on a single microvillus (*cis-*binding). And second, even if CDHR2 were to bend and face CDHR5 on the same microvillus, the geometric alignment is unlikely to be sufficiently precise to permit a simultaneous 2:2 interaction.

We identified various molecular features to answer question (2). First, both CDHR2 and CDHR5 showed a distinctively high thermostability. The *T*m values of CDHR2 and CDHR5 are surprisingly higher than those of other cadherins (CDH17 EC1-4 produced in mammalian cell; 60.1 ℃(30), P-cadherin EC1-2 produced in *E. coli*; 55.4 ℃(31), and PCDH15 produced in *E. coli*; 37.9 ℃(32)). Secondly, the affinity of the *trans*-hetero interaction was among the strongest within the cadherin superfamily(25, 33). Due to this strength, the intermicrovillar adhesion will be maintained robustly under environments with significant shear stress. Another critical property in shear-dominated environments is molecular flexibility, because it provides a mechanism to dissipate those destructive shear forces (Figure 7). The flexibility of EC3–4 domains of CDHR5, connected by a Ca²⁺-free linker, may contribute to force dispersion. As a protease substrate, flexibility leads to the consequence of its quick degradation(34); however, CDHR5 seems to protect itself utilizing a glycan shield to cover it (Figure 6G). In CDHR2, on the contrary, all Ca²⁺-binding motifs between EC repeats are conserved, suggesting that the EC repeats are relatively rigid. Still, like that of PCDH15(35), MAD (membrane adjacent domain) seems to be connected without Ca^2+^, there may be analogous dynamics in this domain. Flexibility may also benefit for the variation of angles among microvilli in the differentiation phase pointed out in recent studies(36, 37). In this way, protocadherin proteins working at microvilli tip have molecular design principles to let hardness and softness coexist in a single molecule.

The kinetics of IMAC formation also depend on the intrinsic affinities of its components; the *K*_D_ value upon the homodimerization of CDHR5 was surprisingly small (i.e. strong affinity) compared to the previously reported *cis*-interaction (38, 39). This strong affinity will decrease the minimum number of molecules required at the tip of microvilli to form *cis*-homodimer, thus facilitating the formation of hetero interaction as well. As a simple estimation, if we suppose that the tip region of a microvillus is a cylinder of 100 nm diameter and 100 nm height, the number of molecules existing there with value of 8 nM is calculated to be less than one molecule, showing that two molecules brought to the tip of microvilli will quickly form *cis*-homodimer to facilitate the IMAC formation.

When CDHR2-CDHR5 interaction is compared with another hetero interacting pair of cadherins, PCDH15-CDH23, expressed in stereocilia of hair cell, we found differences between two systems. In PCDH15-CDH23 interaction called “handshake model”, BSA of the interface is wider than that of CDHR2/CDHR5 (1182.2 Å^2^/interface)(27), and also can happen between monomeric PCDH15 and CDH23(40). Since stereocilia does not have to be closely packed in a hexagonal pattern, less strict control for the establishment of hetero interaction may be necessary. Kinked conformation derived from Ca^2+^-free linker between cadherin EC repeats has been reported also in human PCDH15 EC9-10 and is thought to be important for its mechano-sensing function(28). However, CDHR5 EC3-4 has more acutely bent conformation, suggesting that the level of bending required for tip link of stereocilia and IMAC is different according to their role of mechano-sensing or stable IMAC formation (Supplementary Figure S10).

Collectively, our study indicates that CDHR2 and CDHR5 possess molecular features highly adapted to the circumstances in which intermicrovillar adhesion works. The conclusions of our study advance the understanding of the molecular mechanism of BB assembly, guiding the future design of therapeutic and research-oriented antibodies, as well as ligands modulating their biological function analogously to classical cadherins(41, 42).

## Materials and Methods

### Cloning of CDHR2 and CDHR5 fragments

DNA coding human CDHR2 (Asn21 to Ser1154 corresponding to EC1-9+MAD) or human CDHR5 (Gln26 to Val451 corresponding to EC1-4) were cloned into a pcDNA3.4 vector (Thermo Fisher) containing a Igκ signal peptide (METGLRWLLLVAVLKGVQC or METDTLLLWVLLLWVPGSTGD) in the N-terminal and His_6_tag in the C-terminal of the sequence. For EC1-4 or EC1-2of CDHR2, residues from Asn21 to Phe480 or residues from Asn21 to Pro235 were used, respectively. For EC1-2 of CDHR5, residues from Gln26 toAla230 were used. For the Fc-conjugated construct of CDHR2 EC1-4, His6tag of the EC1-4 construct was replaced with Fc and hinge region of human IgG. Site-directed mutagenesis was performed using KOD -plus-mutagenesis kit (Toyobo) following the instructions of the manufacturer.

### Expression and purification of recombinant proteins

All the CDHR2 or CDHR5 constructs except Fc-conjugation one and CDH17 EC1-4 (Gln23-Asp441) were expressed and purified in the following method. Expi293F cells (Thermo Fisher) were transfected with expression vectors using Gxpress 293 transfection kit (Gmep, Japan) following the instructions of the manufacturer, and supernatant was collected 7 days after transfection. The supernatant was dialyzed against basic buffer (20 mM Tris, 300 mM NaCl, 3 mM CaCl_2_, pH 8.0) supplemented with 5 mM imidazole (binding buffer) and loaded on a cOmplete^TM^ His-tag purification resin (Roche) equilibrated with binding buffer. The resin was washed with basic buffer containing 20 mM imidazole (washing buffer), and subsequently, each protein was eluted with basic buffer containing 300 mM imidazole. Fractions containing the protein of interest were further purified by size exclusion chromatography using a Superdex 200 16/600 GL column (Cytiva) equilibrated with basic buffer.

For Fc-conjugated construct of CDHR2, the culture supernatant was dialyzed against PBS buffer and loaded on Protein A column (Cytiva) equilibrated with PBS buffer. The resin was washed with PBS buffer and subsequently, the protein of interest was eluted with citrate buffer at pH 2.7. Upon elution, the sample was neutralized with 1 M Tris-HCl at pH 9.0 supplemented with 3 mM CaCl_2_. The resultant protein was further purified by size exclusion chromatography using a Superdex 200 16/600 GL column (Cytiva) equilibrated with basic buffer.

Each purified protein was concentrated to about 1.0 mg/mL, frozen in liquid nitrogen, and then stored at -80 ℃. The concentration of each protein was determined from the absorbance at 280 nm measured by a Nanodrop 1000 instrument (Thermo Fisher) and extinction coefficients.

### Cryo-EM sample preparation and data collection

Thawed protein sample of CDHR5 EC1-4 (40 µM) and CDHR2 EC1-4-Fc (40 µM) were mixed and incubated at 4 ℃ for 1 hr. The complex of CDHR2 and CDHR5 was purified by size exclusion chromatography using a Superdex 200 increase 10/300 GL column (Cytiva) equilibrated with assay buffer (10 mM HEPES, 150 mM NaCl, 3 mM CaCl_2_ pH 7.5). The concentration of the complex of CDHR2 and CDHR5 was adjusted to 1 mg/mL, and 3 µL of the solution was applied onto a Quantifoil holey carbon grid (R1.2/1.3, Cu, 200 mesh). Prior to the freezing operation, the grid was glow-discharged at 25 mA for 60 s with GLOQUBE Plus (Quorum Tech.). Excess sample was removed with a blotting time of 6 s and a blotting force of 5 at 4 ℃, 100% humidity before the plunge freezing of the grid in liquid ethane using a Vitrobot Mark IV (Thermo Fisher). A total of 12,275 movies were collected with a K3 detector (Gatan) on a CRYO ARM II 300 (JEM-3300; JEOL) operated at 300keV with an in-column Ω energy filter. Movies were collected in CDS mode using dose-fractionated illumination at a nominal magnification of 60,000x with a physical pixel size of 0.7946 Å/pixel and a defocus range of -1.2 to -2.0 µm for a total dose of 50 e^−^/Å^2^. All movies were automatically acquired with image shift correction using SerialEM(43).

### Cryo-EM data processing and model building

Data processing pipeline was graphically summarized in Supplementary Fig.S2. Movies of the complex of CDHR2 EC1-4-Fc and CDHR5 EC1-4 were imported into cryoSPARC v4.7.0 for motion correction and CTF estimation(44). A subset of all the movies was used to generate 2D averages and *ab initio* reconstitution and then used to produce the template to be subjected to Template Picker. Three consecutive 2D classifications yielded 314,426 particles for further *ab initio* reconstitution and homogeneous refinement. 114,589 particles underwent 3D classification and homogeneous refinement, from which 67,345 particles were used for further 3D classification and non-uniform refinement(45) to produce the final map at 3.41 Å as an overall resolution. The initial model of CDHR2 EC1-4 and CDHR5 EC1-4 from AlphaFold3(46) were fitted into the map using ChimeraX software(47). The resultant model was further refined with a Real-space refinement command of the Phenix software(48). The output structure was further manually refined using the Coot software package(49). Figures were prepared using ChimeraX software. Statistics of data collection and model refinement are summarized in Supplementary Table S2.

### Molecular dynamics (MD) simulations

To investigate the dynamic nature of CDHR2-CDHR5 complex, the MD simulations were performed. As a starting structure, the coordinates of CDHR2 EC1-2 and CDHR5 EC1-4 were extracted from the whole structure determined by cryo-EM. The MD simulations were performed using GROMACS 2015.1(50) with the CHARMM36m force field(51). The glycosylation was modeled using Glycan Reader & Modeler(52, 53). A rectangular box was prepared such that the minimum distance to the edge of the box was 12 Å under the periodic boundary conditions, in which the structures were solvated with TIP3P water(54). The charge of the protein was neutralized by adding Na^+^ or Cl^−^, and the additional ions were added to imitate a salt concentration of 0.15 M of NaCl and 0.003 M of CaCl_2_. Each system was energy-minimized for 5000 steps with the steepest descent algorithm and then equilibrated with the position restraints of protein-heavy atoms and the NVT ensemble, in which the temperature was increased from 50 to 298 K in 150 ps. Following these, simulations with the NVT ensemble for 50 ps and then the NPT ensemble for 200 ps were carried out. During these steps, protein atoms were restrained, while the non-protein atoms moved freely. After all these short simulations, 100 ns of the non-restrained simulations were performed with the NPT ensemble at 298 K. Throughout the simulations, the time step was set to be 2 fs. The cutoff distance of the van der Waals interactions was 12 Å. Long-range electrostatic interactions were evaluated using the particle-mesh Ewald method(55). The covalent bonds involving hydrogen atoms were constrained using the LINCS algorithm(56). A snapshot was saved every 100 ps. Three independent simulations were performed with different initial velocities. The RMSD values or spatial trace were computed using the GROMACS package. The trajectory analysis was performed for those from 25 to 100 ns.

### SEC and SEC-MALS

To investigate the oligomeric state of some cadherin construct, proteins were tested with size exclusion chromatography (SEC). For Figure S1A and Figure 4D, samples adjusted to 25-40 µM were injected into a superdex 200 increase 10/300 column (Cytiva) equilibrated with the assay buffer. To further determine the molecular weight of several cadherin constructs, size exclusion chromatography coupled with a multi-angle light scattering (SEC-MALS) was conducted. For EC1-9+MAD of CDHR2, a Superose 6 increase 10/300 GL column (Cytiva) was used, while for the other samples (CDHR5 EC1-4, EC1-2 and CDHR2 EC1-4-Fc, EC1-4, EC1-2, the complex of CDHR5 EC1-4 and CDHR2 EC1-4-Fc), a superdex 200 increase 10/300 column (Cytiva) equilibrated with the assay buffer was used to obtain the peaks to analyze. The light scattering was detected with DAWN 8+ or DAWN (Wyatt). Each sample was adjusted to 25 to 40 µM and injected into the column stated above. The protein concentration was calculated from UV and the refractive index with a protein conjugate analysis method using dn/dc = 0.185 for the protein portion and dn/dc = 0.145 for the modified glycans. Data were analyzed with ASTRA 8.1.2 or ASTRA 6 software (Wyatt) and molar mass values were determined by the Debye fitting of angle-dependent light scattering.

### Mass Photometry

Determination of dissociation constant of CDHR5 EC1-4 homodimer was performed using the Two^MP^ system (Refeyn) at room temperature. Sample of CDHR5 EC1-4 was dialyzed against the assay buffer and adjusted to 5, 8, 12, 20, 30, 50, 75 nM and incubated for 1 hr at room temperature to let the solution reach equilibrium. The molecular weight and population of each was calculated using DiscoverMP software (Refeyn) with a buffer-free method using Bovine Serum Albumin and Bovine Thyroglobulin (Sigma Aldrich) as a calibration standard. The dissociation constant was determined following equations defined in the previous study(24). Briefly, the dimer fraction was plotted against the total monomer concentration, and the dissociation constant was calculated by performing least-square fitting against the plots.

### Surface plasmon resonance (SPR)

To determine the dissociation constant in hetero interaction (CDHR2-CDHR5), surface plasmon resonance (SPR) was performed using a Biacore 8K instrument (Cytiva). All the samples were dialyzed against the assay buffer supplemented with 0.005% Tween20 and this buffer was directly used as a running buffer. 100 nM of CDHR2 EC1-4-Fc was immobilized on the Sensor Chip Protein A (Cytiva) for 30 s at 10 µL/sec to obtain around 900 RU of immobilization level. Samples of CDHR5 EC1-4 or its mutants were adjusted to the concentration of 1200, 400, 133, 44.4, 14.8, 4.94 nM and injected into the sensor chip surface at a rate of 30 µL/sec. The association time and dissociation time were 60 s or 180 s, respectively. After each cycle, the sensor chip was regenerated by addition of 10 mM Glycine-HCl pH 1.5 for 30 s. The kinetics parameters and dissociation constant were calculated with BIA Evaluation software (Cytiva) using 1:1 binding model.

### Differential scanning calorimetry (DSC)

Thermal stability of each cadherin construct was measured by differential scanning calorimetry (DSC) using a PEAQ-DSC instrument (Malvern). Proteins were dialyzed into the assay buffer and the concentration was adjusted to 1 mg/mL. Measurements were performed at a scan rate of 1℃ per minute from 10℃ to 90℃. Filtered assay buffer from the dialysis was used as the reference sample to obtain the baseline. The melting temperature *T*m was determined by using MicroCal PEAQ-DSC software-in the instrument.

### Differential scanning fluorimetry (DSF)

To cross-validate the DSC results, DSF was performed using a CFX Connect Real-Time System (Bio-Rad). Each sample of cadherin construct at 11 µM in the assay buffer was mixed with SYPRO Orange Protein Gel Stain (50000x concentrated in dimethyl sulfoxide (DMSO); Thermo Fisher) in final concentration of 500x. The protein solution was loaded onto Hard-Shell 96 well PCR Plates (Bio-Rad). Fluorescence emitted from SYPRO Orange upon binding to the exposed hydrophobic surface was monitored to trace the denaturation of each protein. The melting temperature of the protein (*T*m) was calculated based on the derivatives of relative fluorescent unit (RFU) over the derivatives of temperature (-d(RFU)/dT).

### Protein Digestion assay

Protein digestion assay was performed based on the previous study(57). Briefly, PK (10 µM) was incubated with CDHR2, CDHR5, CDH17 (25 µM) for 3 hr at room temperature in the assay buffer. The reaction was quenched by adding 1 µL of PMSF (final concentration = 4 mM). Each sample was tested with SDS-PAGE stained with Coomassie Brilliant Blue (CBB). Band intensities of the CBB-stained SDS-PAGE gel were quantified using ImageJ software (NIH). Images were converted to 8-bit grayscale, and rectangular regions of interest (ROIs) of identical size were applied to each band. Background signal was measured from an adjacent band-free area within the same lane and subtracted from the band integrated density. Relative protein levels were calculated after background correction.

## Acknowledgements

We thank the staff of the Structural Drug Discovery Center via Green Pharma for excellent technical support to obtain cryo-EM data set. Cryo-EM data processing was partially supported by the High-Performance Computing server at CPOS, HKU. We appreciate the technical assistance from The Research Support Center, Research Center for Human Disease Modeling, Kyushu University Graduate School of Medical Sciences, which is partially supported by the Mitsuaki Shiraishi Fund for Basic Medical Research. The supercomputing resources in this study were provided by the Human Genome Center at the Institute of Medical Science, The University of Tokyo, Japan.

## Funding and additional information

This project was funded by grants from the Japan Society for the Promotion of Science (24K18262 to A.S.), The Uehara Memorial Foundation(to A.S.), Koyanagi Foundation (to A.S.), Nakatani Foundation (to A.S.), The Hitachi Global Foundation (to A.S.), the Enhanced New Staff Start-up Research Grant from The University of Hong Kong and 48th PDF/RAP Scheme (to R.H.) and the Platform Project for Supporting Drug Discovery and Life Science Research (Basis for Supporting Innovative Drug Discovery and Life Science Research [BINDS]) from the Japan Agency for Medical Research and Development (grant number JP22ama121033 to K.T., JP22ama121037 to M.K., and JP23ama121031 to S.O., N.M., and J.M.M.C.).

## Author contributions

A.S. and J.M.M.C. conceptualization; A.S., P.A.G.P., K.F., H.K., T.D., K.K., T.T., and J.M.M.C. methodology; A.S., P.A.G.P., K.F., H.K., T.D., K.K., T.T., and J.M.M.C. formal analysis; A.S., P.A.G.P., K.F., H.K., T.D., K.K., T.T., and J.M.M.C. investigation; A.S., K.K., Y.Y., S.Y., M.N., S.N., K.M., S.O., N.M., R.H., K.T. and J.M.M.C. resources; A.S., P.A.G.P., K.F., H.K., T.D., K.K., T.T., and J.M.M.C. data curation; A.S. and J.M.M.C. writing original draft; A.S., P.A.G.P., K.F., H.K., T.D., K.K., T.T., Y.Y., S.Y., M.N., S.N., K.M., S.O., N.M., R.H., K.T., and J.M.M.C. writing-review & editing; A.S. and J.M.M.C. supervision; A.S., Y.Y., S.Y., M.N., S.N., K.M., S.O., N.M., R.H., K.T., and J.M.M.C. project administration; A.S., S.Y., S.O., K.M., R.H., K.T., and J.M.M.C. funding acquisition.

## Data availability

The coordinates of CDHR2-CDHR5 complex have been deposited in the RCSB Protein Data Bank (PDB ID; 24UB). EM density map for the structure has been deposited in the Electron Microscopy Data Bank (Accession code; EMD-69821). The other data are available within the manuscript.

