## Supplementary Information for "Structural basis for the intestinal protocadherin-based intermicrovillar adhesion complex"

### SUPPLEMENTARY TABLES

**Table S1 | Molecular weight of each construct**

| Protein | Calculated Mw (kDa) from *ONE* chain (= A)[a] | Calculated Mw (kDa) from *TWO* chains (= B)[a] | Experimental Mw (kDa) (= C)[b] | \|A-C\|[c] | \|B-C\|[d] | Oligomerization state[e] |
|---|---|---|---|---|---|---|
| CDHR5 EC1-4 in complex with CDHR2 EC1-4-Fc | 158.4[f] | 316.8 | 302 | 143.6 | 14.8 | Hetero-tetramer |
| CDHR5 EC1-4 | 60.7 | 121.4 | 123 | 62.3 | 1.6 | Homodimer |
| CDHR5 EC1-2 | 32 | 64 | 34 | 2 | 20 | Monomer |
| CDHR2 EC1-4 | 70.2 | 140.4 | 82.9 | 12.7 | 57.5 | Monomer |
| CDHR2 EC1-2 | 34.1 | 68.2 | 31.2 | 2.9 | 37 | Monomer |
| CDHR2 EC1-9MAD | 166.8 | 333.6 | 257 | 90.2 | 76.6 | Homodimer |
| CDHR2 EC1-4-Fc | 97.7 | 195.4 | 191 | 93.3 | 4.4 | Homodimer |

a: Mw calculated from polypeptide chain and glycosylation. All putative glycosylation sites based on Uniprot are assumed to be glycosylated. Molecular weight for glycosylation was supposed to be 2 kDa each.

b: The values were determined by SEC-MALS. See Supplementary Figure S1 for raw data.

c: The difference between calculated Mw and experimental Mw in case oligomerization state is monomer.

d: The difference between calculated Mw and experimental Mw in case oligomerization state is dimer.

e: $|A-C| > |B-C|$ means the oligomerization state of the sample is monomer, while $|A-C| < |B-C|$ means it is dimer.

f: In these cases, Mw from a chain of CDHR2 EC1-4-Fc and a chain of CDHR5 EC1-4 is calculated.

**Table S2 | Statistics of data collection and refinement in cryo-EM analysis.**

| | |
|---|---|
| **Data collection** | |
| Equipment | Cryo ARM 300 |
| Voltage (kV) | 300 |
| Sph. Abe. (Å) | 2.7 |
| Camera | K3 |
| Pixel size (Å) | 0.7946 |
| Total Dose (e-/Å$^2$) | 50 |
| Exposure time (s) | 4.19441 |
| Dose rate (e-/Å$^2$/s) | 6.16527 |
| **Refinement** | |
| Chains | 4 |
| Residues | 1772 |
| Water | 0 |
| Ligand | |
| $Ca^{2+}$ | 22 |
| NAG | 44 |
| MAN | 4 |
| Bonds (RMSD) | |
| Length (Å) | 0.003 |
| Angles (°) | 0.541 |
| MolProbity score | 2.01 |
| Clash score | 9.68 |
| Ramachandran plot (%) | |
| Outliers | 0.06 |
| Allowed | 8.39 |
| Favored | 91.55 |
| Rotamer outliers | 0.33 |
| Cβ outliers (%) | 0 |

**Table S3 | Per residue accessible surface area (ASA) and buried surface area (BSA)[a] in CDHR5 homodimerization observed in cryo-EM structure.**

| Chain A | ASA | BSA |
|---|---|---|
| Lys138 | 144.95 | 11.05 |
| Leu232 | 77.73 | 15.77 |
| Thr241 | 112.65 | 25.81 |
| Phe242 | 128.1 | 108.89 |
| Asp244 | 71.09 | 11.88 |
| Tyr246 | 117.82 | 18.7 |
| Val247 | 88.86 | 66.83 |
| Cys248 | 3.2 | 1.4 |
| Ile249 | 103.59 | 102.59 |
| Gln250 | 49.14 | 36.45 |
| Gln252 | 114.21 | 34.56 |
| Phe292 | 96.34 | 12.04 |
| Arg293 | 149.29 | 22.28 |
| Leu324 | 26.11 | 14.56 |
| Arg334 | 76.2 | 17.62 |
| Tyr335 | 120.46 | 63.41 |
| Ser336 | 10.88 | 8.53 |
| Val337 | 62.03 | 61.21 |
| Thr338 | 7.46 | 5.72 |
| Gln339 | 74.27 | 23.55 |
| **Total** | **1634.4** | **662.85** |

| Chain D | ASA | BSA |
|---|---|---|
| Leu232 | 88.94 | 3.18 |
| Pro234 | 54.35 | 1.34 |
| Thr241 | 111.38 | 18.22 |
| Phe242 | 91.65 | 79.88 |
| Ser243 | 99.86 | 9.16 |
| Asp244 | 73.87 | 23.36 |
| Tyr246 | 105.85 | 13.67 |
| Val247 | 84.6 | 69.96 |
| Ile249 | 84.11 | 52.72 |
| Gln250 | 58.63 | 45.56 |
| Gln252 | 100.48 | 41.5 |
| Phe292 | 113.42 | 23.14 |
| Arg293 | 146.18 | 18.73 |
| Leu324 | 27.11 | 21.43 |
| Arg334 | 90.89 | 32.95 |
| Tyr335 | 115.69 | 69.83 |
| Ser336 | 12.7 | 7.15 |
| Val337 | 63.23 | 62.22 |
| Thr338 | 4.01 | 4.01 |
| Gln339 | 69.44 | 34.85 |
| **Total** | **1596.4** | **632.86** |

a: These values were calculated by using PISA server.

**Table S4 | Per residue accessible surface area (ASA) and buried surface area (BSA)[a] in CDHR2 homodimerization observed in cryo-EM structure.**

| Chain | ASA | BSA |
|---|---|---|
| Asn126 | 72.03 | 9.11 |
| Thr127 | 117.41 | 33.47 |
| Ala128 | 66.85 | 18.35 |
| Phe129 | 32.15 | 26.11 |
| Ser130 | 75.41 | 36.7 |
| Ser132 | 77.65 | 0.12 |
| Ile167 | 80.56 | 15.91 |
| Pro168 | 38.14 | 5.19 |
| Tyr193 | 60.54 | 32.02 |
| Ala198 | 32.27 | 21.56 |
| Phe199 | 116.8 | 106.76 |
| Tyr200 | 2.21 | 0.51 |
| Gln201 | 57.23 | 54.89 |
| Pro224 | 64.12 | 6.99 |
| Val225 | 26.34 | 1.34 |
| Phe226 | 91.77 | 62.08 |
| Ser228 | 51.12 | 49.27 |
| Ile229 | 2 | 2 |
| Ser230 | 46.52 | 33.91 |
| Total | 1111.12 | 516.29 |
| Arg243 | 155.39 | 107.86 |
| Glu244 | 158.75 | 74.97 |
| Phe245 | 99.28 | 48.99 |
| Tyr246 | 20.96 | 4.66 |
| Ser247 | 59.68 | 40.01 |
| Gly271 | 35.11 | 2.18 |
| Total | 529.17 | 278.67 |
| Tyr375 | 82.7 | 0.12 |
| Val376 | 7.03 | 5.19 |
| Asp377 | 53.76 | 0.73 |
| Ser381 | 65.9 | 12.92 |
| Pro382 | 63.14 | 3.32 |
| Arg383 | 203.82 | 99.41 |
| Ile384 | 69.27 | 58.17 |
| Pro385 | 74.11 | 32.92 |
| Ile386 | 5.89 | 0.81 |
| Asp387 | 64.1 | 18.7 |
| Total | 689.72 | 232.29 |
| **Total** | **2330** | **1027.3** |

| Chain | ASA | BSA |
|---|---|---|
| Val123 | 91.87 | 0.67 |
| Asn126 | 76.34 | 13.96 |
| Thr127 | 94.99 | 83.26 |
| Ala128 | 70.5 | 46.26 |
| Phe129 | 12.32 | 2.94 |
| Ser130 | 72.13 | 41.33 |
| Ile167 | 66.13 | 12.22 |
| Pro168 | 37.24 | 13.91 |
| Phe199 | 90.2 | 68.23 |
| Gln201 | 63.07 | 61.12 |
| Leu223 | 139.18 | 53.58 |
| Pro224 | 64.55 | 10.36 |
| Val225 | 14.54 | 2.01 |
| Phe226 | 71.71 | 49.83 |
| Ser228 | 40.47 | 38.63 |
| Ile229 | 1.05 | 1.05 |
| Ser230 | 58.74 | 30.54 |
| Total | 1065.03 | 529.9 |
| Gln240 | 118.62 | 36.21 |
| Phe241 | 8.15 | 1.13 |
| Val242 | 91.86 | 61.38 |
| Arg243 | 132.51 | 82.21 |
| Glu244 | 124.88 | 58.04 |
| Phe245 | 128.13 | 34.71 |
| Tyr246 | 11.97 | 0.98 |
| Glu264 | 76.68 | 0.34 |
| Glu264 | 76.68 | 0.34 |
| Total | 692.8 | 275 |
| Ser381 | 75.12 | 4.03 |
| Pro382 | 53.41 | 1.47 |
| Arg383 | 212.39 | 124.93 |
| Ile384 | 79.92 | 65.18 |
| Pro385 | 72.91 | 53.86 |
| Ile386 | 4.32 | 3.22 |
| Asp387 | 87.28 | 21.27 |
| Gln430 | 48.85 | 1.41 |
| Total | 634.2 | 275.37 |
| **Total** | **2392** | **1080.3** |

a: Values were calculated by using PISA server.

*We found three small patches of interfaces in the homodimer of CDHR2. The amino acid residues belonging to each patch are grouped by light orange, orange or brown. See Supplementary Figure S3D for the location of each patch.

**Table S5 | Per residue accessible surface area (ASA) and buried surface area (BSA)[a] in hetero interaction.**

**-Interface between chain A (CDHR5) and chain B (CDHR2)**

**CDHR5**

| Chain A | ASA | BSA |
|---|---|---|
| Gln28 | 139.25 | 16.61 |
| Tyr29 | 115.73 | 59.53 |
| Cys30 | 0.86 | 0.49 |
| Asp35 | 58.27 | 0.37 |
| Ile36 | 101.57 | 55.39 |
| Glu38 | 105.94 | 7.36 |
| Leu65 | 121.57 | 7.2 |
| Tyr86 | 55.46 | 11.16 |
| Ser90 | 60.73 | 32.45 |
| Leu91 | 83.78 | 64.27 |
| Glu93 | 44.02 | 25.58 |
| Thr103 | 63.12 | 5.34 |
| Leu104 | 121.81 | 1.96 |
| Val105 | 69.22 | 39.16 |
| Thr106 | 20.01 | 19.03 |
| Gln107 | 106.92 | 52.16 |
| Leu108 | 36.99 | 18.6 |
| Arg109 | 100.16 | 71.64 |
| Phe111 | 56.27 | 50.04 |
| Ser113 | 35.74 | 29.98 |
| Val114 | 3.48 | 0.12 |
| Leu115 | 64.93 | 9.2 |
| Lys156 | 145.88 | 83.08 |
| Asp157 | 42.86 | 8.36 |
| Asp158 | 87.75 | 8.72 |
| Ile159 | 102.05 | 60.88 |
| Asn181 | 67.64 | 26.61 |
| Arg182 | 104.68 | 5.44 |
| **Total** | **2116.7** | **770.73** |

**CDHR2**

| Chain B | ASA | BSA |
|---|---|---|
| Val22 | 101.04 | 4.18 |
| Ala23 | 39.96 | 24.55 |
| Lys25 | 122.72 | 15.59 |
| Met30 | 20.17 | 1.1 |
| Thr31 | 88.41 | 18.66 |
| Ser32 | 55.34 | 17.13 |
| Ile34 | 114.71 | 31.2 |
| Tyr87 | 41.83 | 28.47 |
| Tyr91 | 118.92 | 77.8 |
| Thr92 | 58.17 | 24.01 |
| Lys94 | 101.94 | 24.03 |
| Ile104 | 70.49 | 22.69 |
| Val106 | 26.28 | 13.05 |
| Gln107 | 82.53 | 0.86 |
| Arg108 | 108.29 | 89.44 |
| Glu109 | 83.38 | 25.87 |
| Leu111 | 55.9 | 45.7 |
| Ile113 | 71.3 | 68.79 |
| Val114 | 4.74 | 1.72 |
| Glu115 | 47.39 | 0.5 |
| Val149 | 92.85 | 1.84 |
| Lys151 | 137.42 | 2.94 |
| Asp152 | 5.64 | 3.55 |
| Met153 | 121.53 | 92.76 |
| Gly154 | 60.86 | 52.87 |
| Ser155 | 58.02 | 33.37 |
| Gly157 | 26.04 | 4.56 |
| Met158 | 105.12 | 81.57 |
| Asn182 | 110.49 | 5.5 |
| Leu209 | 82.07 | 38.48 |
| **Total** | **2072.6** | **824.05** |

**-Interface between chain D (CDHR5) and chain C (CDHR2)**

**CDHR5**

| Chain D | ASA | BSA |
|---|---|---|
| Gln28 | 134.95 | 32.03 |
| Tyr29 | 123.68 | 57.32 |
| Cys30 | 5.1 | 2.7 |
| Ile36 | 99.45 | 46.53 |
| Glu38 | 114.7 | 26.31 |
| Gln58 | 42.23 | 4.51 |
| Leu65 | 110.21 | 10.44 |
| Tyr86 | 65.89 | 12.63 |
| Ser90 | 60.57 | 35.12 |
| Leu91 | 84.48 | 60.1 |
| Glu93 | 28.69 | 9.2 |
| Gln95 | 9.29 | 0.16 |
| Thr103 | 88.6 | 13.89 |
| Val105 | 95.6 | 54.11 |
| Thr106 | 32.41 | 30.46 |
| Gln107 | 95.61 | 50.92 |
| Leu108 | 35.96 | 14.54 |
| Arg109 | 87.85 | 67.8 |
| Phe111 | 74.66 | 65 |
| Ser113 | 32.62 | 29.25 |
| Leu115 | 69.96 | 19.91 |
| Lys156 | 155.14 | 4.35 |
| Ile159 | 90.43 | 7.36 |
| Asn181 | 66.57 | 0.16 |
| Arg182 | 115.98 | 1.17 |
| **Total** | **1920.6** | **655.97** |

**CDHR2**

| Chain C | ASA | BSA |
|---|---|---|
| Lys25 | 118.96 | 8.42 |
| Thr31 | 84.86 | 30.93 |
| Ser32 | 46.6 | 21.58 |
| Ile34 | 107.68 | 27.13 |
| Tyr87 | 46.48 | 30.54 |
| Tyr91 | 117.43 | 71.46 |
| Thr92 | 56.14 | 20.52 |
| Lys94 | 112.15 | 44.92 |
| Tyr103 | 106.24 | 0.24 |
| Ile104 | 78.28 | 1.34 |
| Arg108 | 81.42 | 0.51 |
| Glu109 | 58.96 | 7.24 |
| Leu111 | 55.1 | 44.85 |
| Ile113 | 69.55 | 67.37 |
| Val114 | 2.48 | 0.73 |
| Glu115 | 53.93 | 0.17 |
| Leu147 | 91.09 | 13.57 |
| Asp152 | 8.75 | 2.94 |
| Met153 | 118.99 | 92.47 |
| Gly154 | 62.98 | 57.12 |
| Ser155 | 60.76 | 37.43 |
| Gly157 | 18.07 | 7.36 |
| Met158 | 116.74 | 99.26 |
| Val159 | 33.21 | 14.55 |
| Leu209 | 86.65 | 44.2 |
| **Total** | **1793.5** | **746.85** |

a: These values were calculated by using PISA server.

**Table S6 | Melting temperatures of CDHR5 mutants measured by differential scanning fluorimetry (DSF).**

| | Peak 1 (℃) | Peak 2 (℃) |
|---|---|---|
| WT | 63.2 | 72.2 |
| S90A | 63.6 | 71.8 |
| L91W | 63.2 | 72.0 |
| Q107A | 63.4 | 72.2 |
| R109G | 62.8 | 71.4 |
| F242R | - | 71.8 |
| V247T | 63.6 | 70.2 |
| Y335D | - | 71.2 |
| V337T | - | 71.8 |

**Supplementary Figures**

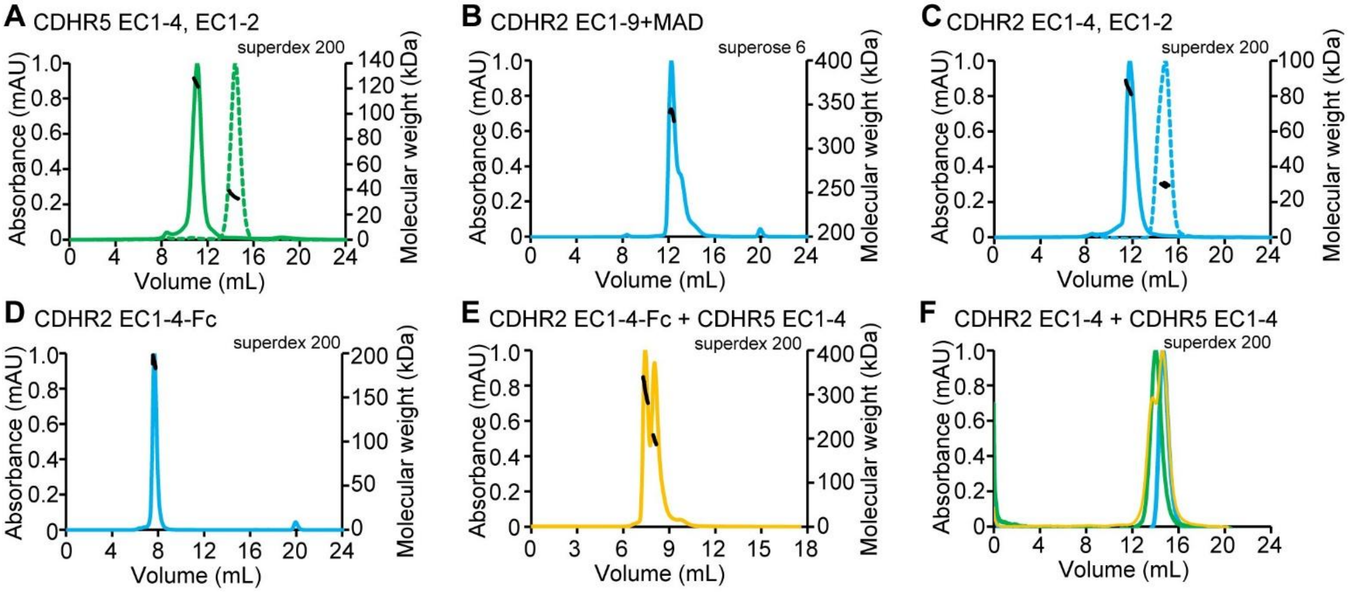


**Figure S1 | Molecular weight determination of cadherin constructs by SEC-MALS. (A)~(E)** SEC-MALS experiments for CDHR5 EC1-4, EC1-2 **(A)**, CDHR2 EC1-9+MAD **(B)**, CDHR2 EC1-4, EC1-2 **(C)**, CDHR2 EC1-4-Fc **(D)** and the mixture of CDHR2 EC1-4-Fc and CDHR5 EC1-4 **(E)**. In **(B)** and **(C)**, results for EC1-4 are shown in solid lines, while those for EC1-2 are shown in dotted lines. In each panel, the molecular weight corresponding to the peak is shown in black traces. Also, the column used for each sample is also written. Panel **(A)** clearly showed that CDHR5 EC1-4 is homodimer, while EC1-2 is monomer. Also, panel **(B)** showed CDHR2 was homodimer in full ectodomains; however, panel **(C)** showed that only EC1-4 or EC1-2 was not sufficient to keep the dimeric state. Therefore, we prepared forced-dimerization construct in which Fc of human IgG was conjugated into CDHR2 EC1-4. Panel **(D)** showed that this construct formed homodimer as expected. In **(E)**, the former peak corresponds to the complex of CDHR2 EC1-4-Fc and CDHR5 EC1-4, while the latter peak mainly corresponds to the unbound CDHR2 EC1-4-Fc. See also Table S1 for the determined molecular weights. **(F)** SEC chromatograms of CDHR2 EC1-4 (blue, notice that this is not EC1-4-Fc), CDHR5 EC1-4 (green) and mixture of CDHR2 EC1-4 and CDHR5 EC1-4 (yellow). The mixture of both EC1-4, which is monomer, did not cause a peak corresponding to their complex. From these results, the hetero interaction is strong only when both CDHR2 and CDHR5 are homodimers, while that between monomer CDHR2 or CDHR5 is too weak to be observed in the SEC experiments with the tested concentrations (~40 μM).

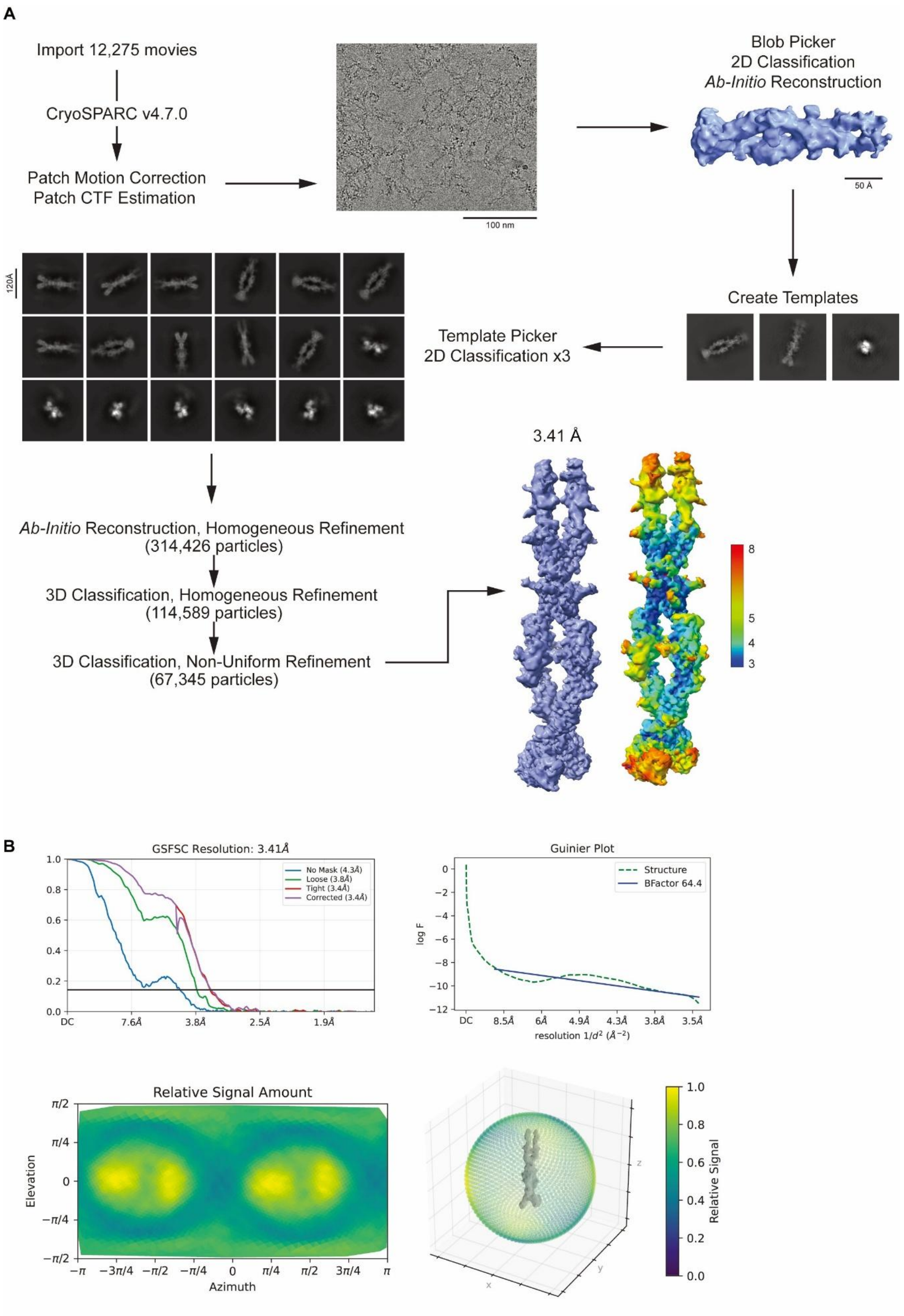

A
Import 12,275 movies
CryoSPARC v4.7.0
Patch Motion Correction
Patch CTF Estimation
100 nm
Blob Picker
2D Classification
Ab-Initio Reconstruction
50 Å
Create Templates
Template Picker
2D Classification x3
120Å
Ab-Initio Reconstruction, Homogeneous Refinement
(314,426 particles)
3D Classification, Homogeneous Refinement
(114,589 particles)
3D Classification, Non-Uniform Refinement
(67,345 particles)
3.41 Å
8
5
4
3
B
GSFSC Resolution: 3.41Å
No Mask (4.3Å)
Loose (3.8Å)
Tight (3.4Å)
Corrected (3.4Å)
DC
7.6Å
3.8Å
2.5Å
1.9Å
Guinier Plot
Structure
BFactor 64.4
log F
8.5Å
6Å
4.9Å
4.3Å
3.8Å
3.5Å
resolution 1/d² (Å⁻²)
Relative Signal Amount
Elevation
Azimuth
Relative Signal

**Figure S2 | Analysis workflow in Cryo-EM. (A)** Schematic showing the cryo-EM data processing workflow. A representative motion-corrected cryo-EM micrograph obtained from a CRYO ARM 300 II instrument, *ab initio* 3D model, 2D classifications, final map and resolution distributions are shown. **(B)** Gold-standard Fourier shell correlation (GSFSC) curve for final full-particle map, Guinier Plot, Relative signal amount from all directions is shown. All are computed with cryoSPARC.

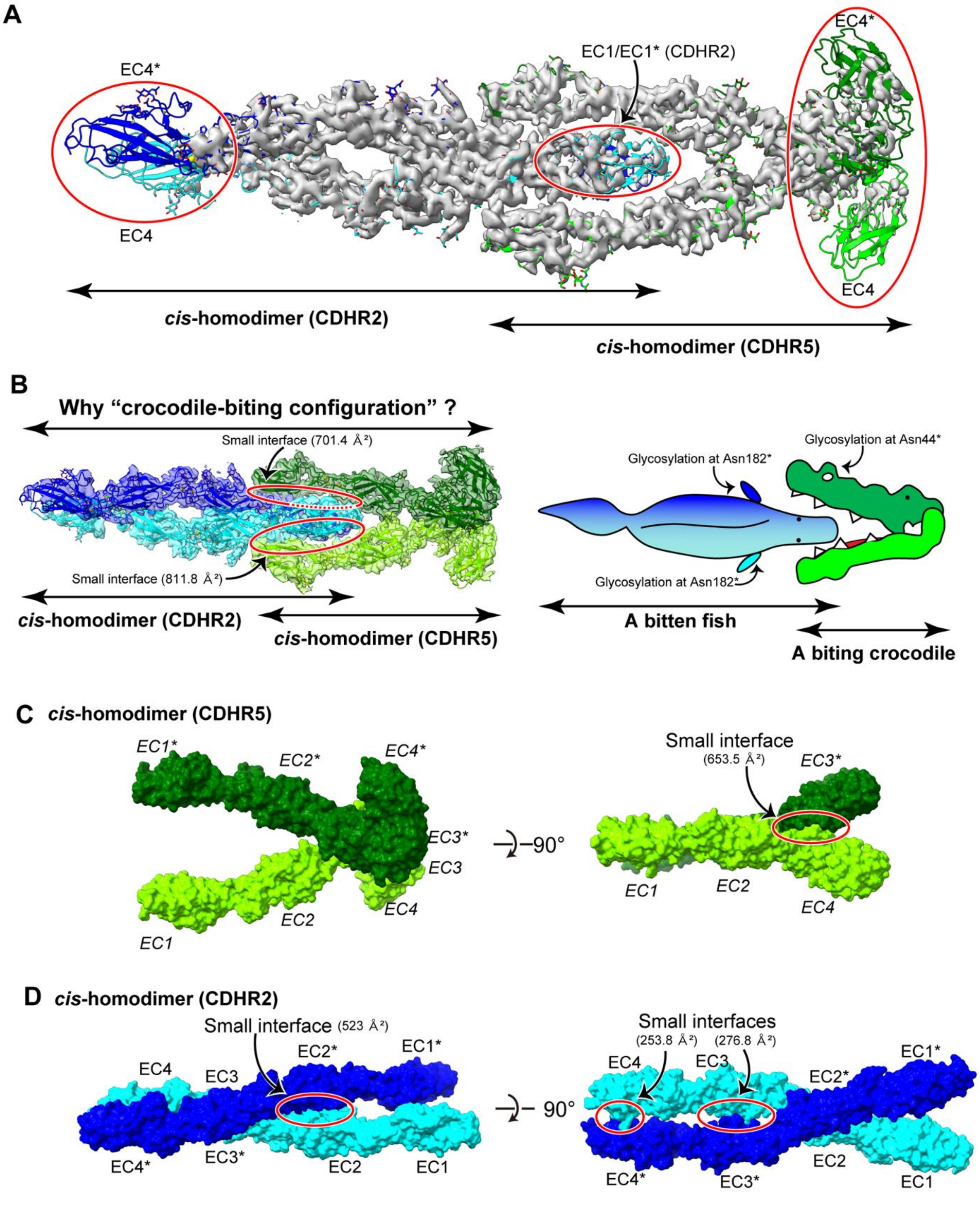


**Figure S3 | Molecular architecture of CDHR2 and CDHR5. (A)** Cryo-EM map with modeled structure shown in cartoons. Chains of CDHR2 are colored in cyan or blue, while those of CDHR5 in light green or deep green. The counter level of the map is increased compared to that shown in Figure 1B to show that EC4/EC4* from CDHR2 and CDHR5 had weak density. **(B)** We named the silhouette of heterotetramer "crocodile-biting configuration". Homodimer of CDHR5 looks like a crocodile opening its mouth widely, representing the EC1-2/EC1-2*. On the other hand, homodimer of CDHR2 looks like a fish about to be bitten by the crocodile. Indeed, glycosylation at Asn182 in CDHR2 and that at Asn44 in CDHR5 added a flavor of fins of the fish and nose of the crocodile, respectively. The interfaces are emphasized by red circles with buried surface area (BSA) written. **(C)** Structure of *cis*-homodimer of CDHR5 extracted from complex structure with surface representation in two angles. The interface is shown in a red circle with BSA value. **(D)** Structure of *cis*-homodimer of CDHR2 extracted from complex structure with surface representation in two angles. *Cis*-homodimer of CDHR2 had three small interfaces; one is at EC2/EC2*, another at EC3/EC3* and the other EC4/EC4*. Similarly to **(C)**, the homo interfaces are shown in red circles with BSA value.

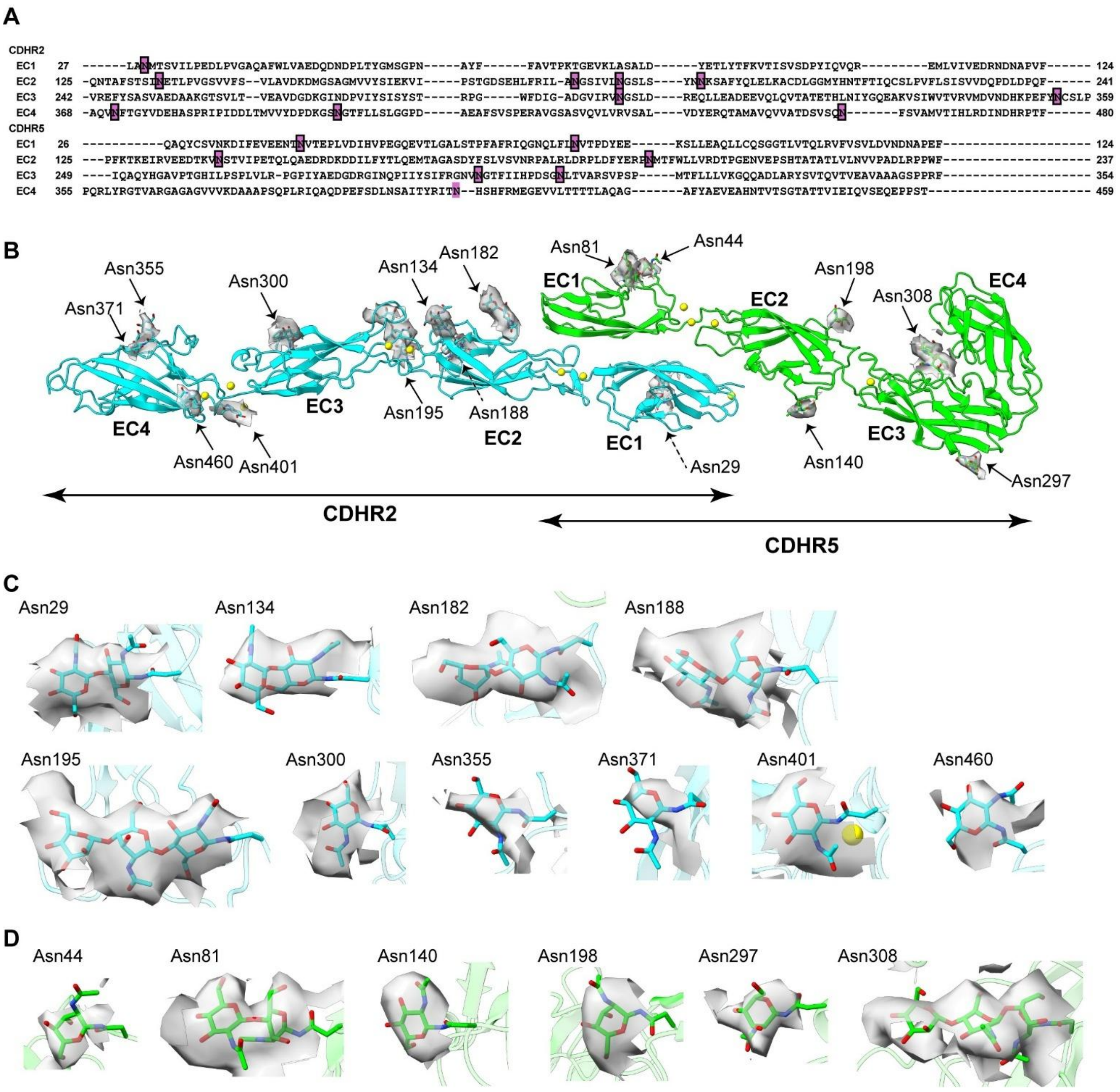


**Figure S4 | Glycosylation of CDHR2 and CDHR5. (A)** Primary sequences of CDHR2 and CDHR5, with putative glycosylation sites purple in their background. Some electron density corresponding to the glycosylation was found in the positions with frame lines, while a position without it did not show clear electron density to model glycans. The numbers on the right or left of each line show the amino acid residue numbering. **(B)** Positions of glycosylation sites from chain A of CDHR5 and chain C of CDHR2. The dotted lines show the glycans in the positions are placed on the back side of the sheet. **(C) (D)** Cryo-EM density corresponding to each glycosylation in CDHR2 **(C)** or CDHR5 **(D)**. The density is cropped until around 3 Å of NAG and (or ) MAN for the glycan.

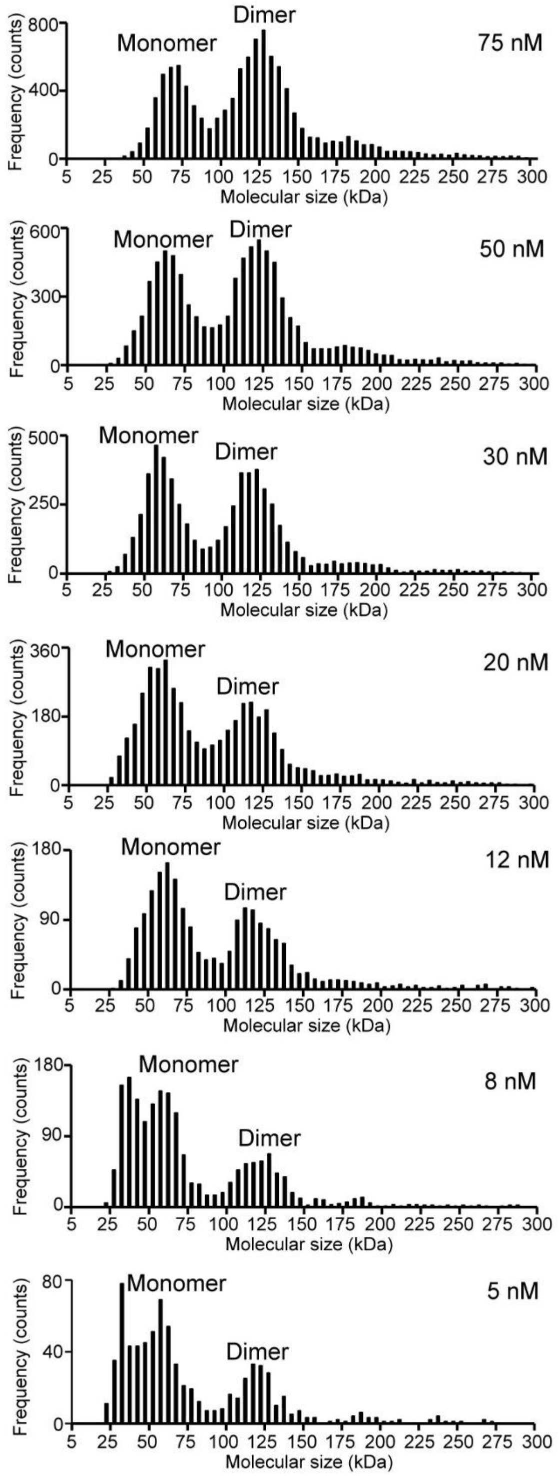


**Figure S5 | Raw data from mass photometry.** The histograms show the distribution of the molecular sizes detected by mass photometry in analyzing CDHR5 EC1-4 at each concentration. The number of counts from each peak corresponding to monomer and homodimer were used to calculate the monomer/dimer fractions.

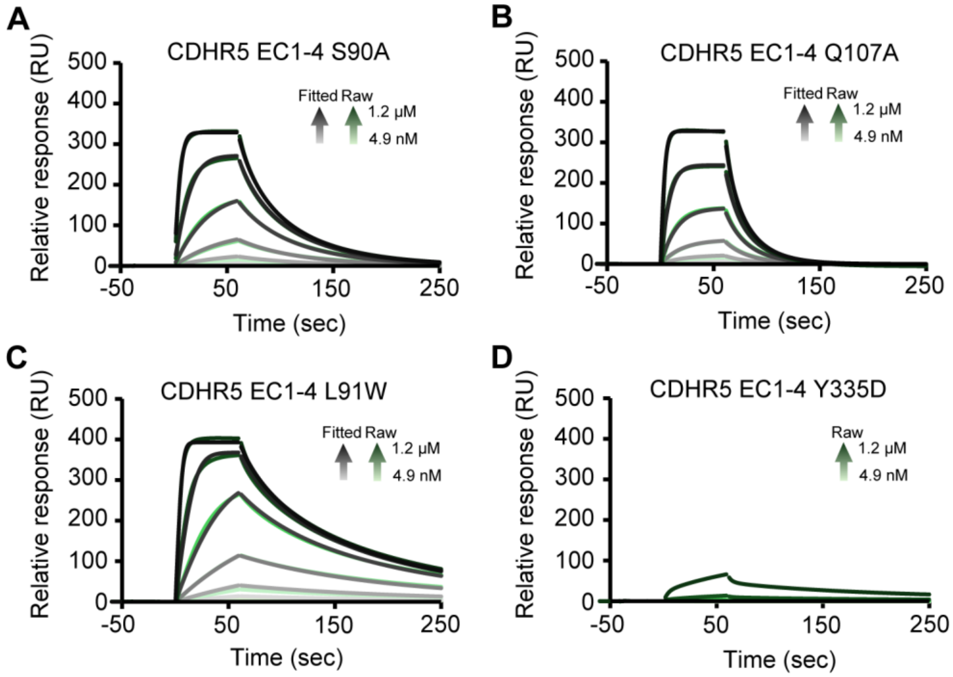


**Figure S6 | Mutational analysis of hetero interaction by SPR. (A)~(D)** SPR sensorgrams of CDHR5 EC1-4 S90A **(A)**, EC1-4 Q107A **(B),** EC1-4 L91W **(C)** and monomer fraction of EC1-4 Y335D **(D)** as an analyte. In all conditions, CDHR2 EC1-4-Fc was immobilized on Sensor Chip Protein A. The analyte solutions were adjusted to 1200, 400, 133, 44.4, 14.8 and 4.94 nM. The green traces show the raw data, while the black traces show the fitted data. For **(D)**, no fitting is shown due to poor binding.

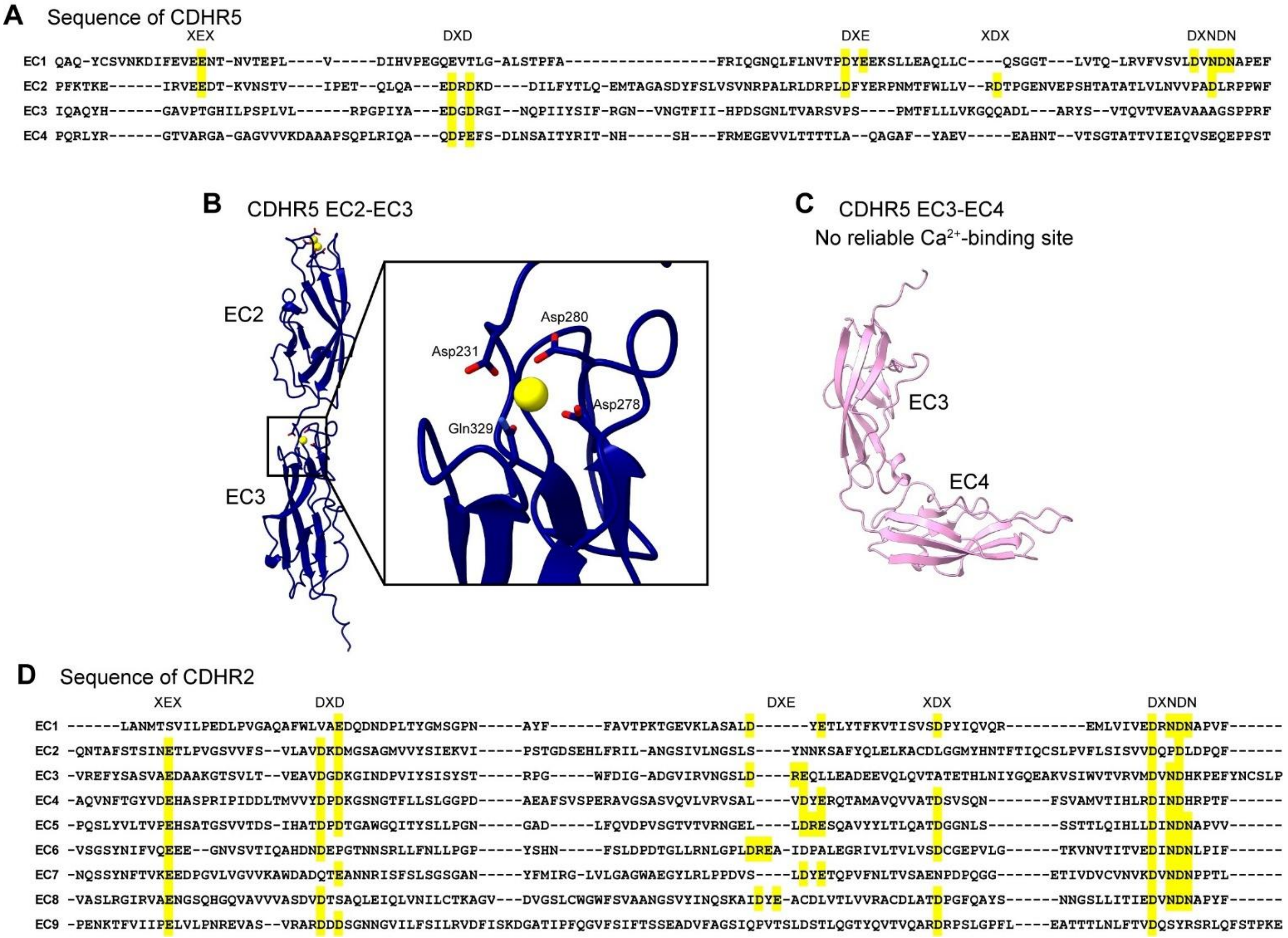


**Figure S7 | $Ca^{2+}$-binding in CDHR5 and CDHR2. (A)** Primary sequence of CDHR5 EC1-4. The canonical $Ca^{2+}$-binding motifs are highlighted in yellow background. This is basically the same as Figure 4A but is shown here for the easy comparison with panel **(D)**. **(B) (C)** Prediction of $Ca^{2+}$-binding to EC2-EC3 **(B)** or EC3-4 **(C)** of CDHR5 by AlphaFold3. In EC2-3, AlphaFold3 predicted only one $Ca^{2+}$ binds to the region, while there was no $Ca^{2+}$-binding to EC3-4 domain. $Ca^{2+}$ with reliability of pIDDT>50 is shown. **(D)** Primary sequence of CDHR2 EC1-9. The canonical $Ca^{2+}$-binding motifs are highlighted in yellow background.

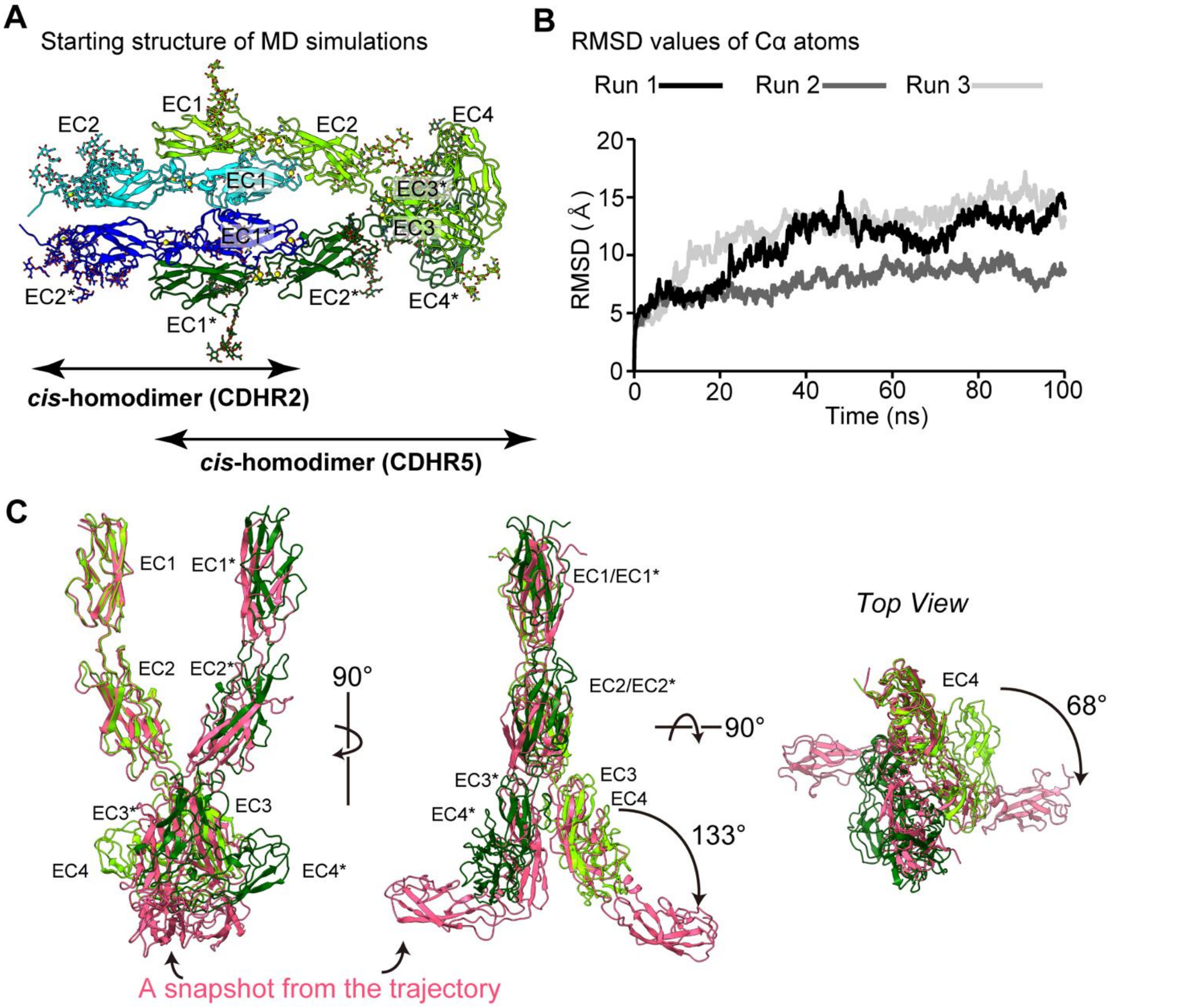


**Figure S8 | MD simulations of CDHR2-CDHR5 complex structure. (A)** Starting structure. EC1-2/EC1-2* from CDHR2 and EC1-4/EC1-4* from CDHR5 were extracted from the whole structure in Figure 1B. Also, N-glycans (G0) were modelled based on CHARMM Glycan Reader and Modeler to the position where electron density of a part of glycosylation was observed from cryo-EM. **(B)** RMSD of Cα atoms from the starting structure. Black, grey and light grey lines show three independent runs performed in MD simulations. **(C)** Superposition of the starting structure (chain A colored in light green, chain B colored in dark green) and a snapshot from the MD trajectories (both chains colored in pink). Because of the flexible nature of EC3-4 domain, the angle between the mass center of EC3 and a Cα atom of a residue in the linker and mass center of EC4 becomes as large as 133° from original angle of 55°. Also, seen from the top view, the twisting movement was also observed shown by a black arrow.

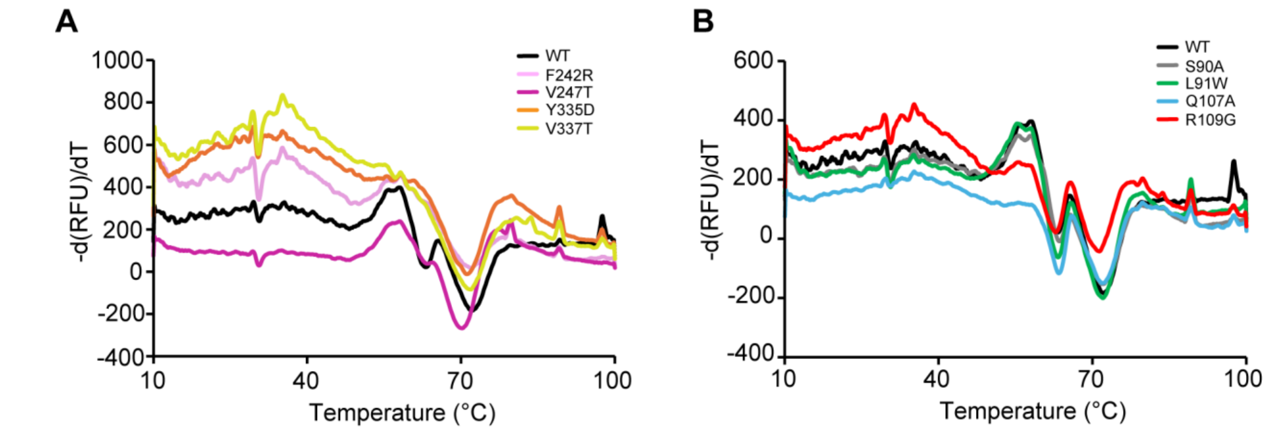


**Figure S9 | Thermal stability of CDHR5 mutants measured by differential scanning fluorimetry (DSF). (A) (B)** Derivatives of relative fluorescence unit (RFU) over the derivatives of temperature (d(RFU)/dT) are shown for mutants tested in analyses of homodimerization **(A)** or of hetero interaction **(B)**. For each panel color codes for mutants are shown.

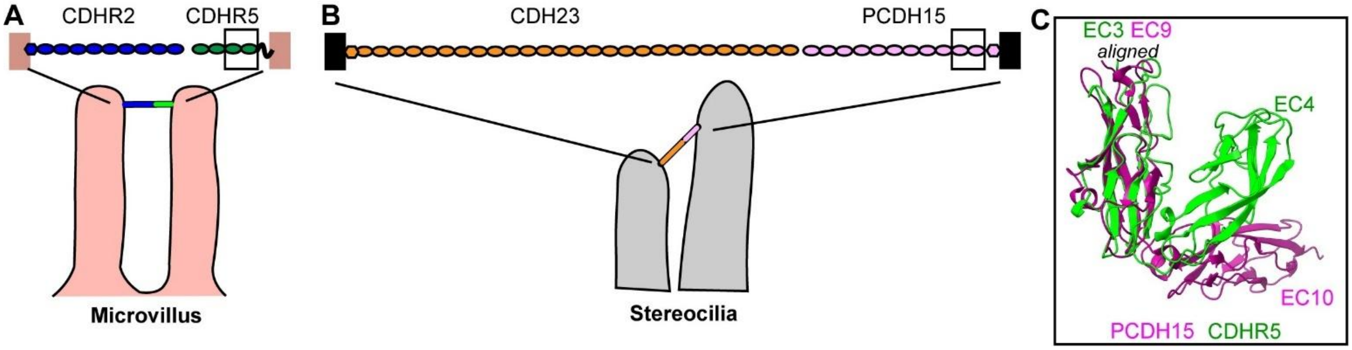


**Figure S10 | Kinked domains observed in PCDH15 and CDHR5. (A), (B)** Schematic illustration of IMAC **(A)**, and stereocilia in hair cells, in which PCDH15 and CDH23 form hetero interaction **(B)**. The black boxes in each panel show the location of domains superposed in panel **(C)**. **(C)** Superposition of CDHR5 EC3-4 (green) and PCDH15 EC9-10 (magenta). EC3 of CDHR5 and EC9 of PCDH15 are used to align both structures. EC3-4 of CDHR5 has more acute inter-domain angle.
